# PBRM1 regulates the stress response in epithelial cells

**DOI:** 10.1101/408575

**Authors:** Elizabeth G. Porter, Alisha Dhiman, Basudev Chowdhury, Benjamin C. Carter, Hang Lin, Jane C. Stewart, Majid Kazemian, Michael K. Wendt, Emily C. Dykhuizen

**Author notes:** These authors contributed equally to this work (Elizabeth G. Porter, Alisha Dhiman).

## Abstract

Polybromo1 (PBRM1) is a chromatin remodeler subunit highly mutated in cancer, particularly renal clear cell carcinoma. PBRM1 is a member of the SWI/SNF subcomplex, PBAF (PBRM1-Brg1/Brm Associated Factors) and is characterized by six tandem bromodomains. Here we establish a role for PBRM1 in epithelial cell maintenance through the expression of genes involved in cell adhesion, metabolism, stress response, and apoptosis. In support of a general role for PBRM1 in stress response and apoptosis, we observe that loss of PBRM1 results in an increase in reactive oxygen species generation and a decrease in cellular viability under stress conditions. We find that loss of PBRM1 promotes cell growth under favorable conditions but is required for cell survival under conditions of cellular stress.

## Introduction

*PBRM1*, a gene which encodes a subunit of the PBAF chromatin remodeling complex, is mutated in over 3% of all cancers with the highest mutation rate occurring in clear cell renal cell carcinoma (ccRCC), where it is mutated in 40-50% of patients (Cancer Genome Atlas Research Network, 2013; Peña-Llopis et al., 2012; Varela et al., 2011). The PBAF chromatin remodeling complex is a minor subcomplex of the human SWI/SNF, or BAF, chromatin remodeling family, subunits of which (*SMARCA4* (BRG1*), ARID1A*, and *SMARCB1 (*SNF5 or BAF47)) are also frequently mutated in cancers (Kadoch et al., 2013; Shain and Pollack, 2013). Along with PBRM1, the PBAF subcomplex exclusively contains ARID2, BRD7, BAF45A, as well as several subunits shared with the more abundant BAF complex (Kaeser et al., 2008; Tatarskiy et al., 2017; Xue et al., 2000). PBRM1 is comprised of several domains associated with binding to chromatin including six tandem bromodomains (BDs), two bromo-adjacent homology (BAH) domains and a high mobility group (HMG), implicating PBRM1 as a chromatin targeting subunit of PBAF. For the most part, the chromatin signatures bound by PBRM1 have not yet been determined, although histone 3 lysine 14 acetylation (H3K14Ac) has been defined as a primary target for the second bromodomain (BD2) *in vitro* (Charlop-Powers et al., 2010), and validated as the acetylation mark most critical for association of the full PBAF complex to histone peptides (Porter and Dykhuizen, 2017). PBRM1 has homology to RSC1/2 and RSC4 subunits of the yeast RSC chromatin remodeling complex, which also interacts with H3K14Ac, particularly during DNA damage (Duan and Smerdon, 2014; Y. Wang et al., 2012). However, unlike subunits of RSC, PBRM1 does not seem to be necessary for viability in the majority of mammalian cell types, and in fact, while PBRM1 is essential for embryonic heart development in mice (X. Huang et al., 2008; Z. Wang et al., 2004), adult mice with knockout of PBRM1 are phenotypically normal except for an age-related hematopoietic stem cell (HSC) defect (Lee et al., 2016).

The most well-defined cellular role for PBRM1 is in DNA-damage repair (Brownlee et al., 2014; Kakarougkas et al., 2014), which is in line with observation of H3K14Ac at sites of DNA damage (Lee et al., 2010); however, the low mutational burden and relative genome stability of PBRM1-mutant tumors makes it unclear as to how a role in DNA-damage repair relates to the tumor suppressive phenotypes of PBRM1(Sato et al., 2013). As such, most of the focus has been on deciphering how transcriptional functions for PBRM1 relate to a role in tumor suppression. Transcriptional profiling of human ccRCC indicate that PBRM1 mutant tumors have a hypoxic transcriptional signature (Sato et al., 2013), which is in agreement with recent reports that PBRM1 mutation amplifies the hypoxia inducible factor (HIF) transcriptional program signature induced upon von Hippel-Lindau (VHL) deletion in cell culture (Gao et al., 2017) and in a mouse model of renal cancer (Nargund et al., 2017). Recent work with kidney specific (KSP and PAX8) Cre mouse models indicate that VHL knockout or PBRM1 knockout alone is not sufficient for cancer formation but that both are required for kidney tumor formation in mice (Espana-Agusti et al., 2017; Gu et al., 2017; Nargund et al., 2017).

While these recent mouse studies have solidified a role for PBRM1 as a bona fide tumor suppressor in renal cancer, the molecular mechanism by which PBRM1 acts as a tumor suppressor is still unclear. For example, PBRM1 exhibits tumor suppressive phenotypes in a subset of cancer cell lines (Chowdhury et al., 2016; L. Huang et al., n.d.; Xia et al., 2008), but PBRM1 knockdown in many cell lines produces no phenotype (Chowdhury et al., 2016; Gao et al., 2017) or even decreases cellular viability (Lee et al., 2016). In the renal cancer setting, this context-specific function is mediated, in part, through HIF1a expression, which is required for PBRM1’s tumor suppressor phenotype in renal cell lines (Murakami et al., 2017) (Shen et al., 2011); however, the context-dependent function observed in other cell types is still undefined. Here we used epithelial cell lines to define how the function of PBRM1 in non-transformed cells may relate to its function as a tumor suppressor. Through genome-wide transcriptional analysis, we have defined a general role for PBRM1 in regulating the expression of genes involved in stress response, particularly endoplasmic reticulum stress and apoptosis. To support this general function, we have found that loss of PBRM1 results in accumulation of reactive oxygen species (ROS) and a failure to induce apoptosis under a variety of-high stress conditions. Based on our findings, we propose that PBRM1 acts to regulate stress response genes that restrain cellular proliferation under low stress conditions but protect cells under high stress conditions.

## Results

### Knockdown of PBRM1 in normal epithelium promotes growth and a loss of epithelial cell maintenance

Since mutation of PBRM1 in epithelial cells is an early event in tumorigenesis (Gerlinger et al., 2014) we set out to understand the tumor suppressive role PBRM1 plays in the context of normal epithelial cells. We depleted *PBRM1* using lentiviral shRNA in several epithelial cell lines including the immortalized human kidney epithelial cell line HK2, the canine kidney epithelial cell line MDCK, and the mouse mammary epithelial cell line NMuMG (SI1A). In addition to its role in renal cancer, PBRM1 acts as a tumor suppressor in mammary epithelium derived cancers as observed in PBRM1 mutated (Xia et al., 2008) and PBRM1-downregulated breast cancers (SI1B) (Mo et al., 2015). The loss of PBRM1 resulted in an increase in proliferation in all of these cell lines (Fig 1A) consistent with a role for PBRM1 as a tumor suppressor in epithelial cells. Since NMuMG is the most commonly used epithelial cell line model, we used it for further analysis of PBRM1’s role in epithelial maintenance. In NMuMG cells, knockdown of PBRM1 decreases protein levels of E-cadherin, a marker of epithelial cells, and increases vimentin, a marker of mesenchymal cells (Fig 1B) (Kalluri and Weinberg, 2009). A decrease in E-cadherin at adherens junctions results in a weakening of cell-cell adhesion and also results in the release of bound β-catenin, which normally anchors E-cadherin to the actin cytoskeleton (Kalluri and Weinberg, 2009). In agreement with this, we observed an increase in nuclear β-catenin signaling upon PBRM1 knockdown (Fig 1C). The complete loss of E-cadherin expression and cellular morphology characteristic of a robust epithelial-to-mesenchymal transition (EMT) was not observed; instead, the observed phenotypes reflect a partial EMT and a reduction in epithelial maintenance. The same phenotypes were also observed upon knockdown of BRD7, another PBAF-specific subunit,(Kaeser et al., 2008) although these findings are complicated by a decrease in PBRM1 upon BRD7 knockdown (SI1C). Consistent with the documented role for the BAF complex in maintaining human mammary epithelial proliferation (Cohet et al., 2010), knockdown of BAF subunits ARID1A or BRG1 resulted in cell arrest and death (data not shown).

**Figure 1:**
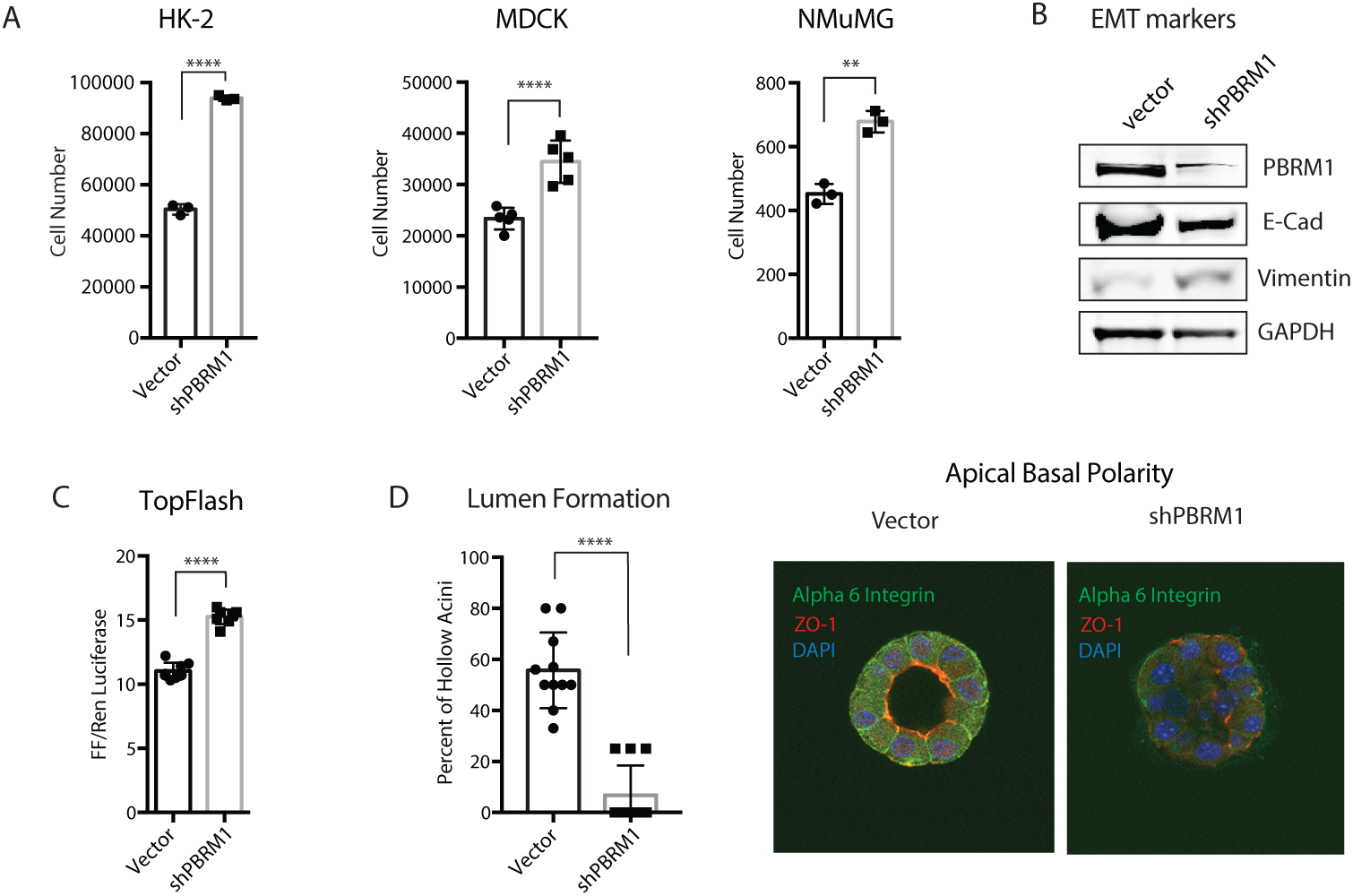
Knockdown of PBRM1 in epithelial cells promotes cell growth and partial epithelial-mesenchymal transition. **A.** Epithelial cell lines HK-2 (human kidney), MDCK (canine kidney) and NMuMG (mouse mammary), were counted after 72h growth and cell number are presented as mean ± SD. n = 3-5. **B.** Immunoblot analysis of whole cell lysates from NMuMG cells indicates that PBRM1 knockdown results in decreased E-cadherin expression and increased vimentin expression. **C.** β-catenin signaling, as measured using Top Flash reporter assay in NMuMG cells with vector control or shPBRM1. Individual replicates are presented as mean ± SD n = 8. **D**. Acini with hollow lumen from NMuMG cells grown in 3D culture for 10 days were counted in a blinded manner and the frequency was calculated from total acini in a field of image (average of 5-6 acini per field). The data from eleven independent images were statistically analyzed (Student t-test) and presented as mean ± SD n = 11. Representative image of acini grown for 14 days were analyzed using immunofluorescence staining with anti-ZO1 (red) and anti-alpha-6-integrin (green). Nuclei (blue) were visualized by DAPI. A designation of *=p <0.05, ** = p <0.01, *** = p < 0.001, **** = p < 0.0001 (paired Student’s t test). ns, not significant. Error bars represent S.D.

A decrease in E-cadherin in epithelial cells during EMT also results in a decrease in cellular polarity, a feature central to epithelial function. To investigate the contribution of PBRM1 to the maintenance of epithelial cell polarity we plated NMuMG cells in Matrigel-based 3D culture where they self-assemble into luminal structures consisting of hollow acini displaying apical-basal polarity (Hall et al., 1982). Upon PBRM1 knockdown, the spheres fail to establish hollow lumen and lose both ZO-1 at apical tight junction and basal/lateral staining of alpha 6 integrin (Figure 1D, SI1D). This phenotype is consistent with that observed in NMuMG epithelial cells with PTEN deletion or PI3K activating mutation (Berglund et al., 2012).

### PBRM1 regulates genes involved in cell adhesion, signaling, stress response, and apoptosis

To identify genes regulated by PBRM1 in epithelial cells, we performed RNA-Seq from control and PBRM1 knockdown NMuMG cells. In total, we identified 2467 genes with significantly increased transcript levels and 1927 genes with significantly decreased transcript levels upon PBRM1 knockdown. Gene ontology (GO) analysis identified numerous pathways significantly enriched in genes downregulated upon shPBRM1, including cell movement, cell structure, development, and signaling (Fig 2A), as would be expected based on the phenotypes observed for PBRM1 in Figure 1. In addition, there were numerous enriched biological pathways involved in stress response, cellular homeostasis, translational elongation, and apoptosis. In contrast, there were few significantly enriched biological pathways for genes upregulated upon shPBRM1, however, these pathways included microtubule-based processes (Fig 2B).

**Figure 2:**
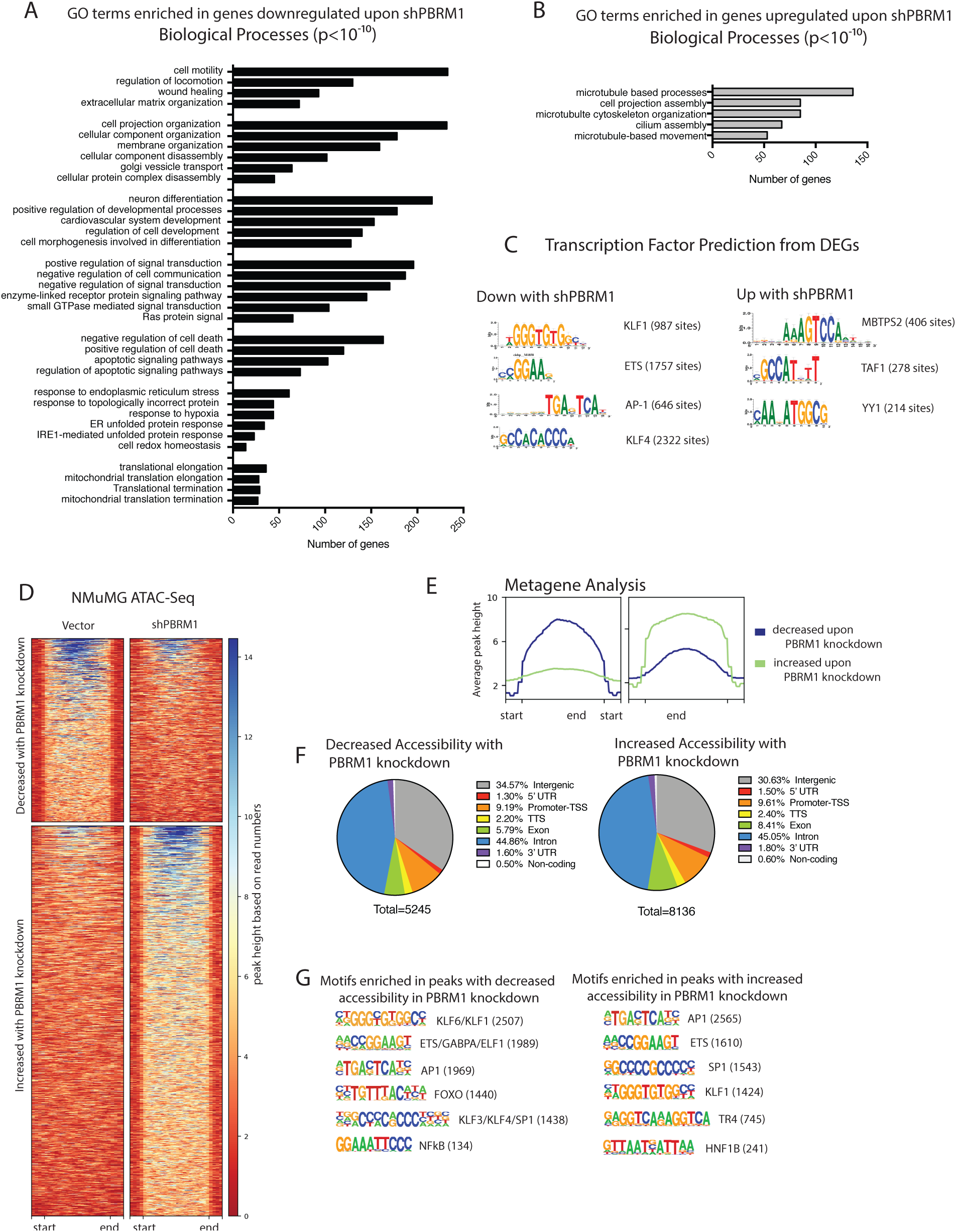
Transcriptional analysis indicates a role for PBRM1 in upregulating genes involved in cell adhesion and migration, response to stress, and apoptosis. **A)** Top overrepresented biological process GO terms (p values<10^-10^) for differentially expressed genes that are downregulated in NMuMG cells with shPBRM1. **B)** Top overrepresented biological process GO terms (p values<10^-10^) for differentially expressed genes that are upregulated in NMuMG cells with shPBRM1. **C)** Putative transcription factors were identified for genes exhibiting differential expression in NMuMG cells with shPBRM1. **D)** Heat maps of regions identified as differentially accessible with PBRM1 knockdown using ATAC-Seq analysis of NMuMG cells. Regions of at least 1.5-fold differential accessibility were calculated between pooled samples of three biological replicates. **E)** Metagene plots of the regions identified as differentially accessible with PBRM1 knockdown using ATAC-Seq analysis of NMuMG cells. **F)** Genomic elements associated with the differentially accessible peaks. The overall distribution was calculated as a percentage of the total differentially accessible regions for each condition. **G)** Motif analysis was performed using HOMER for the differentially accessible peaks. Statistically significant motifs were identified based on relative enrichment over genomic areas with similar AT content.

### PBRM1 is predicted to cooperate with transcription factors involved in response to stress

We further utilized the RNA-Seq datasets to predict upstream regulators that might cooperate with PBRM1 in transcription of target genes. For genes downregulated upon shPBRM1, several enriched consensus sequences were identified, with the most robust identified for KLF transcription factors, including KLF4, which is required for epithelial cell homeostasis (Ghaleb et al., 2011; Yu et al., 2012) such that the KLF4 knockdown in NMuMG cells has similar phenotypes to the PBRM1 knockdown (Tiwari et al., 2013). In addition there was significant enrichment for AP-1 transcription factors, such as JUN/FOS, which are upregulated during stress (Jiang et al., 2016; Rössler and Thiel, 2017), as well as ATF4 and ATF6, which are key transcriptional regulators in ER stress response (de Nadal et al., 2011). Similar to the RNA-Seq analysis, very few TF consensus sequences were enriched in the promoters of genes upregulated upon shPBRM1, but the main consensus sequence enriched were associated with MBTPS2, a protease that activates transcription factors involved in cholesterol synthesis and ER stress response (Rawson, 2013), and YY1, a structural protein involved in promoter-enhancer associations (Beagan et al., 2017).

To further define whether these putative transcription factors are directly regulated by PBRM1’s chromatin remodeling function, we next turned to ATAC-Seq to identify sites of PBRM1-dependent chromatin accessibility. As observed elsewhere (Gao et al., 2017), PBRM1 knockdown did not have dramatic effects on global chromatin accessibility. It did, however, result in a significant decrease in accessibility (at least 1.5-fold) at 5,245 sites and increased accessibility at 6,790 in NMuMG cells (Fig 2D, E) with similar genomic distributions (Fig 2F). Similar results were obtained using PBRM1 knockdown in HK2 epithelial cells (SI2). To identify transcription factors (TFs) that are potentially dependent on PBRM1 for chromatin binding, we calculated the enrichment of TF consensus binding sequences at sites with differential accessibility upon PBRM1 knockdown (Fig 2G). Several TF consensus sequences were significantly enriched compared to background at sites of increased as well as at sites of decreased accessibility upon shPBRM1. Consensus sequences for KLF, AP-1, ETS, FOXO, and NFkB transcription factors were highly enriched in regions with decreased accessibility upon shPBRM1, which correlates with the predicted regulators based on RNA-seq data (Fig 2D). In addition, there was a significant overlap between genes downregulated upon shPBRM1 and genes with an associated region of decreased accessibility (536 genes, p = 4.5×10^-83^) and these regions displayed enrichment for KLF, AP-1, ETS, and FOXO consensus sequences. While we observed similar enrichment of consensus sequence binding sites in the regions with increased accessibility upon shPBRM1, the regions of accessibility didn’t correlate to genes upregulated upon shPBRM (144 genes, p = 0.455). In summary, a unifying role in stress response was apparent for the putative TFs identified for both PBRM1-dependent gene activation and PBRM1-dependent chromatin accessibility.

### Knockdown of PBRM1 results in elevated ROS generation under cellular stress conditions

Due to the transcriptional signature indicating an increased importance for PBRM1 in regulating genes involved in stress response, we next examined how depletion of PBRM1 affects reactive oxygen species (ROS), which is generated by cells under a variety of cellular stresses (Geou-Yarh Liou, 2010). NMuMG cells have low endogenous ROS levels and PBRM1 knockdown results in a small but significant increase in ROS under normal cell culture conditions (Fig 3A), as measured by conversion of 2′,7′-dichlorodihydrofluorescein diacetate (H_2_DCFDA) to the highly fluorescent 2′,7′-dichlorofluorescein (DCF) by intracellular ROS. To understand the effects of PBRM1 on ROS under high cellular stress, we looked at ROS levels after recovery from hydrogen peroxide treatment (Fig 3B), glucose deprivation (Fig 3C), hypoxia-inducing CoCl2 treatment (Fig 3D), and DNA-damaging doxorubicin treatment (Fig 3E). Under all of these stress conditions, cells lacking PBRM1 expression displayed higher amounts of ROS.

**Figure 3:**
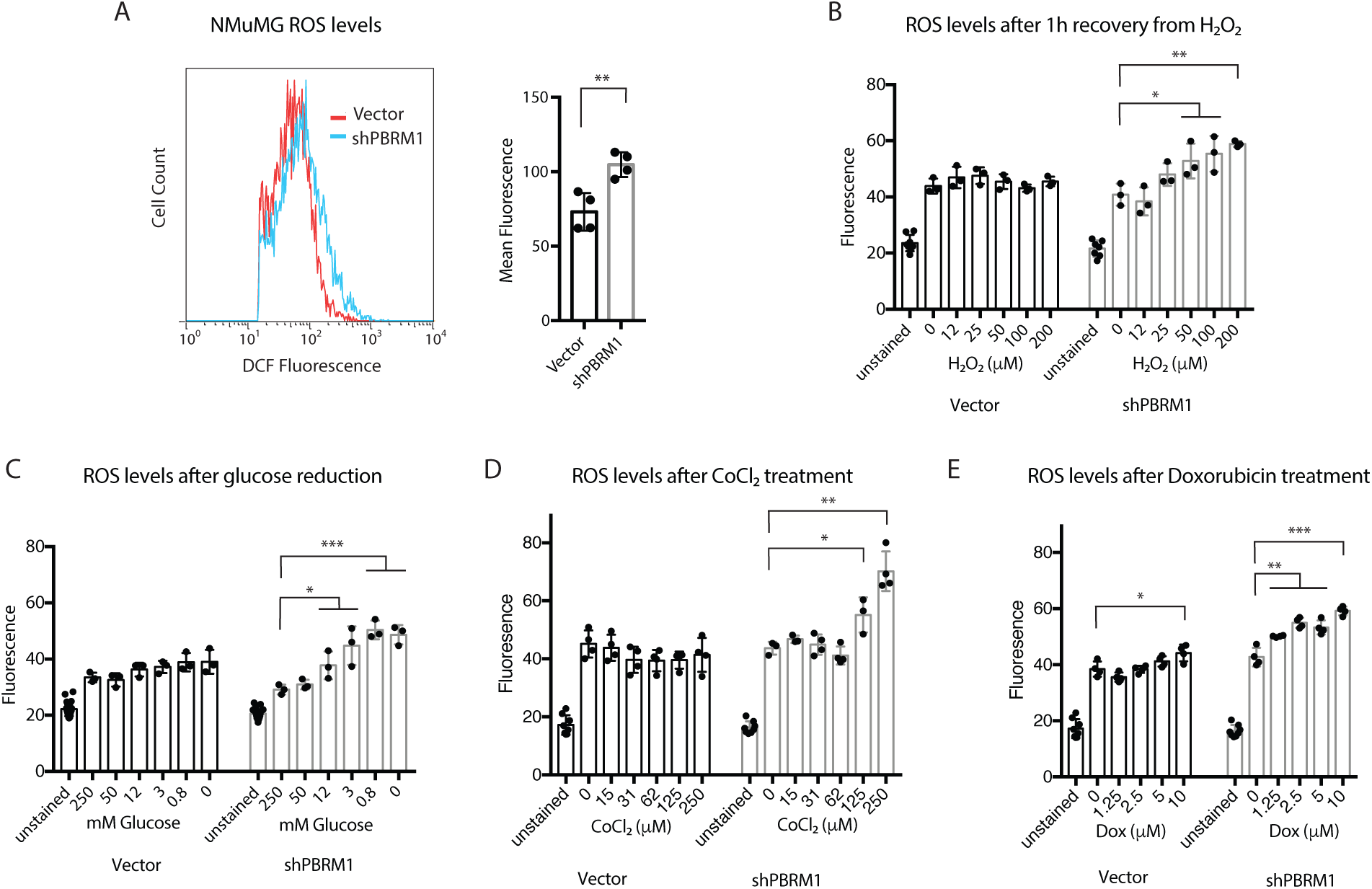
Loss of PBRM1 results in increased reactive oxygen species under cellular stress conditions. **A**) NMuMG cells were trypsinized and stained for 30 min with H_2_-DCFDA, washed with PBS, and analyzed using flow cytometry. The mean fluorescence value for 10,000 cells was calculated from four independent experiments. **B**) NMuMG cells grown in 96-well plates were treated with increasing concentrations of H_2_O_2_ for 1 h, washed with PBS, and incubated with H_2_-DCFDA for 30 min. Reagent was washed away and the DCF fluorescence was measured in live cells. **C**) NMuMG cells grown in 96-well plates were treated with media containing varying concentrations of glucose (normal media = 250 mM) for 16 h, washed with PBS, and incubated with H_2_-DCFDA for 30 min. Reagent was washed away and the DCF fluorescence was measured in live cells. **D**) NMuMG cells grown in 96-well plates were treated with media containing varying concentrations of CoCl_2_ for 24 h, washed with PBS, and incubated with H_2_-DCFDA for 30 min. Reagent was washed away and the DCF fluorescence was measured in live cells. **E**) NMuMG cells grown in 96-well plates were treated with media containing varying concentrations of doxorubicin for 24h, washed with PBS, and incubated with H_2_-DCFDA for 30 min. Reagent was washed away and the DCF fluorescence was measured in live cells. A designation of *=p <0.05, ** = p <0.01, *** = p < 0.001, **** = p < 0.0001 (paired Student’s t test). ns, not significant. Error bars represent S.D.

### PBRM1 expression is cytoprotective under high stress conditions

A low level increase in ROS production promotes cancer progression by increasing signaling, and facilitating transformation through increasing genomic instability and inflammation (Fig 4A)(Geou-Yarh Liou, 2010). In addition, ROS can increase AKT phosphorylation and induce changes in cell adhesion molecules to increase motility (Geou-Yarh Liou, 2010), both of which we observed previously in Caki2 renal cancer cells without PBRM1 (Chowdhury et al., 2016) and in epithelial cells lacking PBRM1 (SI 1A). Further, the increase in proliferation upon PBRM1 knockdown in NMuMG cells can be reversed upon addition of exogenous antioxidant Vitamin C, supporting an oncogenic role for low level increases in ROS (Fig 4B). While increases in ROS are characteristic in cancer and contribute to transformation and oncogenesis, cancer cells need to avoid extremely high levels of ROS due to cytotoxicity (Fig 4A) (Geou-Yarh Liou, 2010). To understand how PBRM1-regulated ROS levels under high stress conditions affect cellular viability, we measured cell survival after high concentrations of hydrogen peroxide for 16h. We found that PBRM1 knockdown decreased viability under these high stress conditions in the NMuMG (Fig 4C) and MDCK epithelial cells (SI3). This was not due to increased apoptosis in the PBRM1 knockdown, and in fact, cells lacking PBRM1 displayed a deficiency in Annexin V (Fig 4D) and cleaved PARP (Fig 4E) under stress conditions. This is in line with the transcriptional role for PBRM1 in regulating pro-apoptotic genes (Fig 2A). While it seems counterintuitive that PBRM1-expressing cells have both increased apoptosis and increased cell survival under high stress conditions, it is consistent with a role for PBRM1 in the stress response, which often results in apoptosis if cellular stresses are not resolved. In contrast, cells lacking PBRM1 are unable to mount a proper response to external stress, leading to high ROS levels and cell death through other means, such as necrosis (Fulda et al., 2010), which is supported by the increase in necrosis marker lactate dehydrogenase (LDH) in the NMuMG cells lacking PBRM1 (Fig 4F).

**Figure 4:**
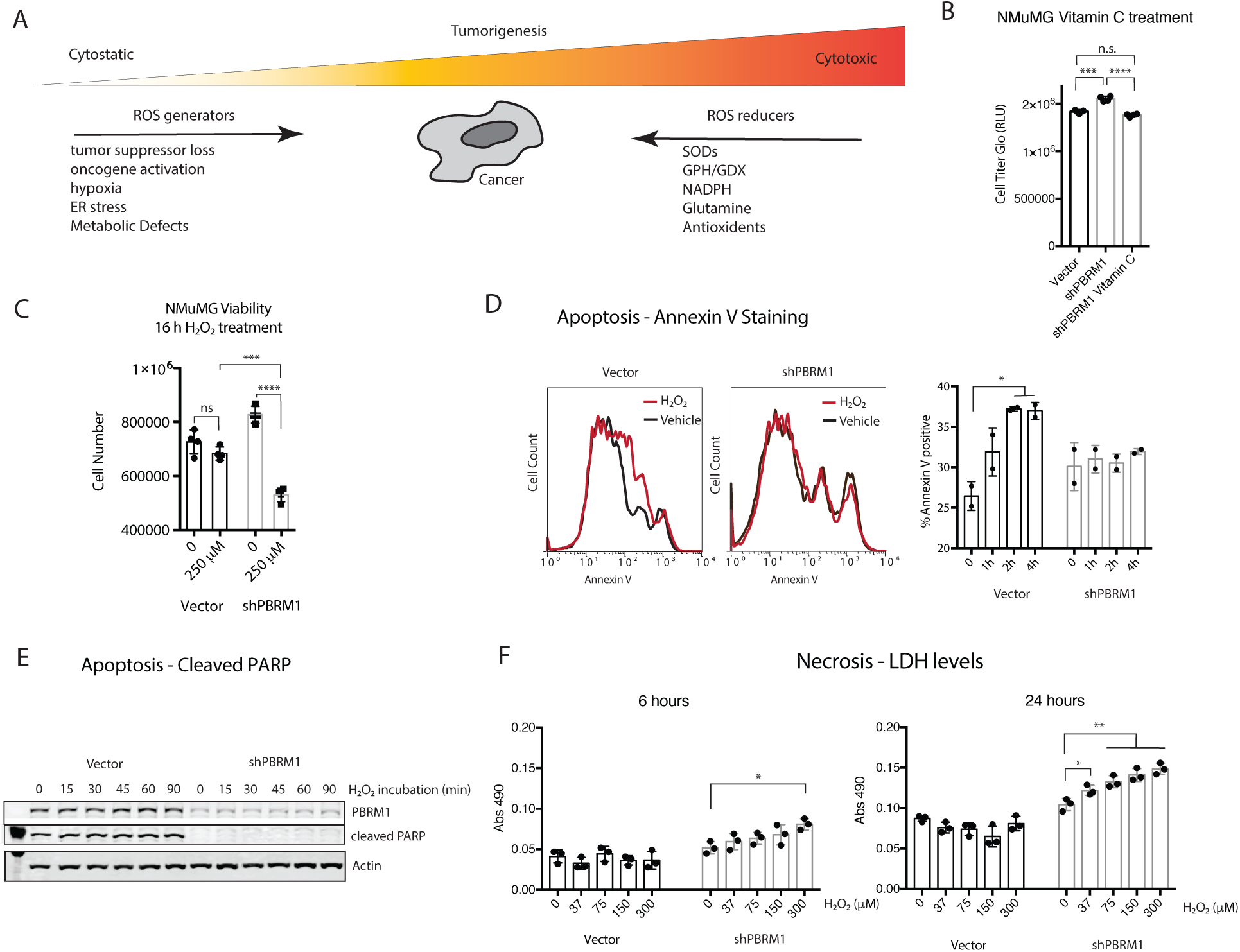
PBRM1 induces apoptotic pathways and increases cellular viability under high cellular stress conditions. **A**) Depiction of the multifaceted role ROS regulators play in cancer **B**)NMuMG cells were cultured for three days in normal media or media supplemented with 20 *μ*g Vitamin C. **C)** NMuMG cells were cultured in normal cell media or 250 *μ*M H_2_O_2_ for 16 h and the number of live cells were counted using trypan blue. **D**) Annexin V staining was measured in NMuMG cells treated with 200 *μ*M H_2_O_2_ for the indicated times. The percentage of Annexin V positive cells was calculated for two independent experiments. **E**) Whole cell lysates were prepared from NMuMG cells treated with 200 *μ*M H_2_O_2_ for the indicated times and probed for the indicated proteins using immunoblot analysis. **F**) LDH release was measured from 10 *μ*L of media using LDH Cytotoxicity Assay Kit II (Abcam). A designation of *=p <0.05, ** = p <0.01, *** = p < 0.001, **** = p < 0.0001 (paired Student’s t test). ns, not significant. Error bars represent S.D.

### PBRM1-regulated transcriptional effects under cellular stress conditions

To determine if the dependency on PBRM1 expression for viability under stress conditions is due to the PBRM1’s regulation of different genes under stress conditions, we characterized the transcriptional profile of NMuMG cells with shPBRM1 grown in H_2_O_2_ (200 *μ*M) for 2h and low glucose media for 6h (Fig 5A). There were between 1000-2000 DEGs identified in cells grown in H_2_O_2_ or low glucose growth conditions for both vector control and shPBRM1 cells (SI4B), although the impact of PBRM1 expression on overall gene expression was more significant than the impact of either stress condition on overall gene expression (SI4A). We observed a significant correlation between gene expression changes induced by the alternative growth conditions in the two cell lines (vector and shPBRM1), indicating that at least some of the gene expression changes are independent of PBRM1 expression (SI4B). In addition, we observed a significant correlation between gene expression changes induced by each growth condition, implying that similar transcriptional alterations are induced by both stress conditions (SI4C). To ask whether different sets of genes were regulated by PBRM1 under high stress conditions compared to low stress conditions, we evaluated the correlation between the differential expression of genes by PBRM1 in both conditions (Fig 5B). In general, the correlation was high between PBRM1-regulated genes under low stress and the PBRM1-regulated genes under high stress conditions, with the highest correlation between PBRM1 regulated genes in the two stress conditions (SI4D). This indicates that while there is some alteration in PBRM1-regulation of gene expression under high stress conditions, most of the gene regulation remains the same. As expected, the significantly enriched GO terms are similar for genes regulated by PBRM1 with stress treatment compared to untreated cells (Fig 5C), just with more genes from pathways involved in cell adhesion, signaling and apoptosis altered by shPBRM1 under stress treatments (Fig 5C). Further, we observed several GO terms that were significantly enriched only for genes downregulated by shPBRM1 under stress treatment, including cell cycle, protein metabolic processes, and cellular response to stress, and for genes downregulated by shPBRM1 only under H_2_O_2_ treatment, such as RNA processing and DNA-damage response (Fig 5D). In conclusion, the RNA-Seq analysis of PBRM1-dependent gene expression under stress conditions supports a role for PBRM1 in suppressing cell cycle and metabolic processes to promote cell survival under conditions of high stress.

**Figure 5:**
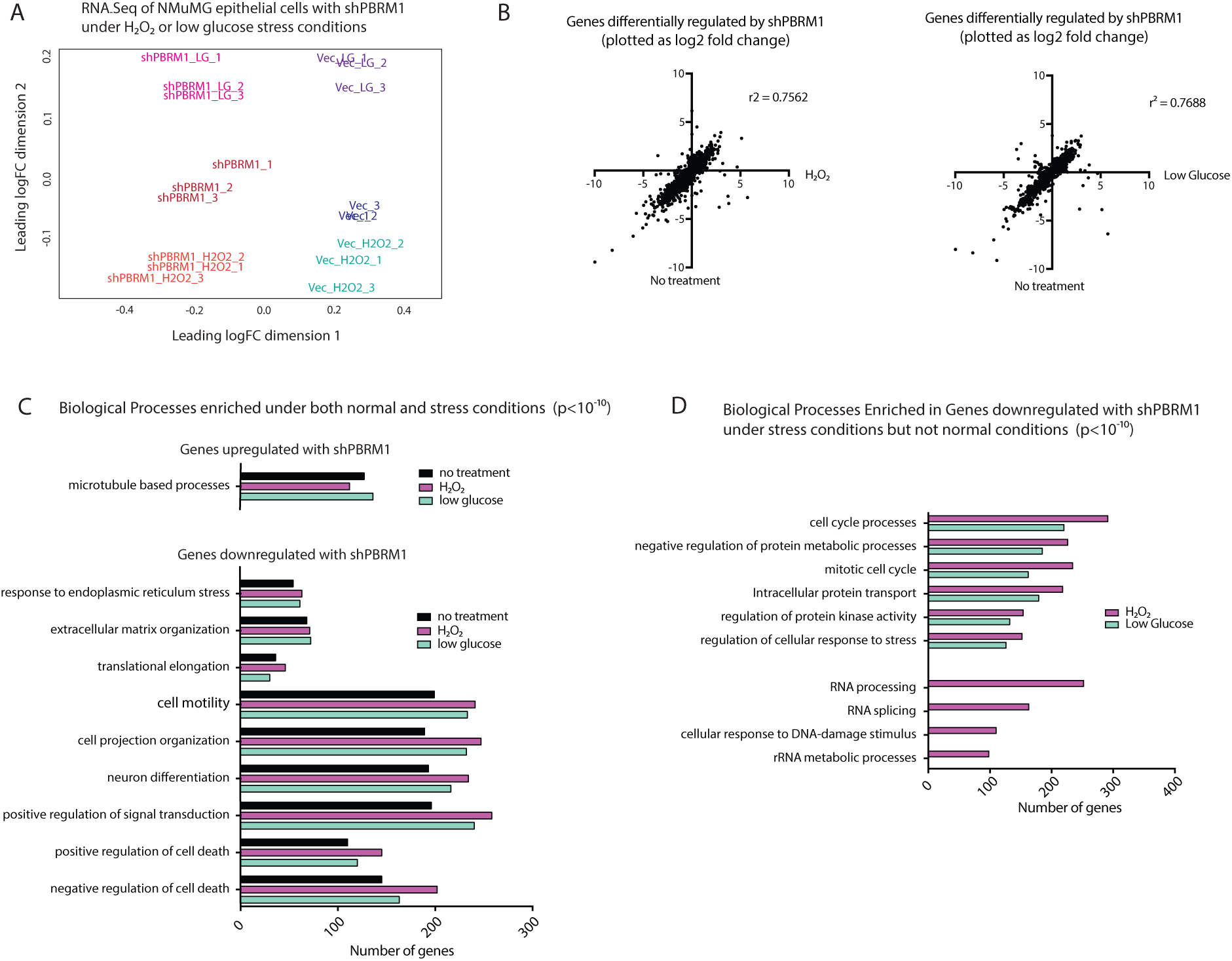
PBRM1 transcriptional regulation is amplified under conditions of cellular stress. **A)** RNA-Seq was performed on NMuMG cells grown in 200 *μ*M H_2_O_2_ for 2h or glucose-free media for 6h. **B**) Each data point represents a single gene, *x axis*: differential expression in H_2_O_2_ treated cells (left) or low glucose treated cells (right) upon shPBRM1, *y axis*: differential expression in normal cell culture conditions upon shPBRM1. The degree of correlation was calculated using all differentially expressed genes. **C)** Top overrepresented biological process GO terms (p values<10^-10^) for differentially expressed genes in NMuMG cells with shPBRM1. **C)** Top overrepresented biological process GO terms (p < 10^-10^) for genes differentially expressed upon shPBRM1 in all cell culture conditions. **D)** Overrepresented biological process GO terms (p < 10^-10^) for differentially expressed genes downregulated upon shPBRM1 only under H_2_O_2_ or low glucose cell culture conditions.

### PBRM1 has cell type specific roles on viability

Since establishing that PBRM1 knockdown can have different effects on viability depending on the stress environment, we re-evaluated the premature senescence phenotype previously described for PBRM1 knockout in mouse embryonic fibroblasts (MEFs) (Lee et al., 2016). We confirmed that PBRM1 knockdown (SI5A) results in a loss in the proliferative capacity of MEFs (Fig 6A) similar to published findings with the PBRM1 conditional knockout (Lee et al., 2016). We next observed a significant increase in ROS levels and H_2_O_2_ levels in MEFs upon PBRM1 knockdown (Fig 6B, SI5B), which was most similar to the robust increase in ROS levels observed in shPBRM1 NMuMG cells grown under high stress conditions. This particular sensitivity of MEFs to PBRM1 knockdown is most likely due to the unique susceptibility of MEFs to oxidative stress from high oxygen content in air (Espana-Agusti et al., 2017). To support this, we found that exogenous antioxidants such as Vitamin C (Fig 6C) or N acetyl cysteine (NAC) (SI5C) were able to reverse the viability defect induced by PBRM1 knockdown in MEFs.

**Figure 6:**
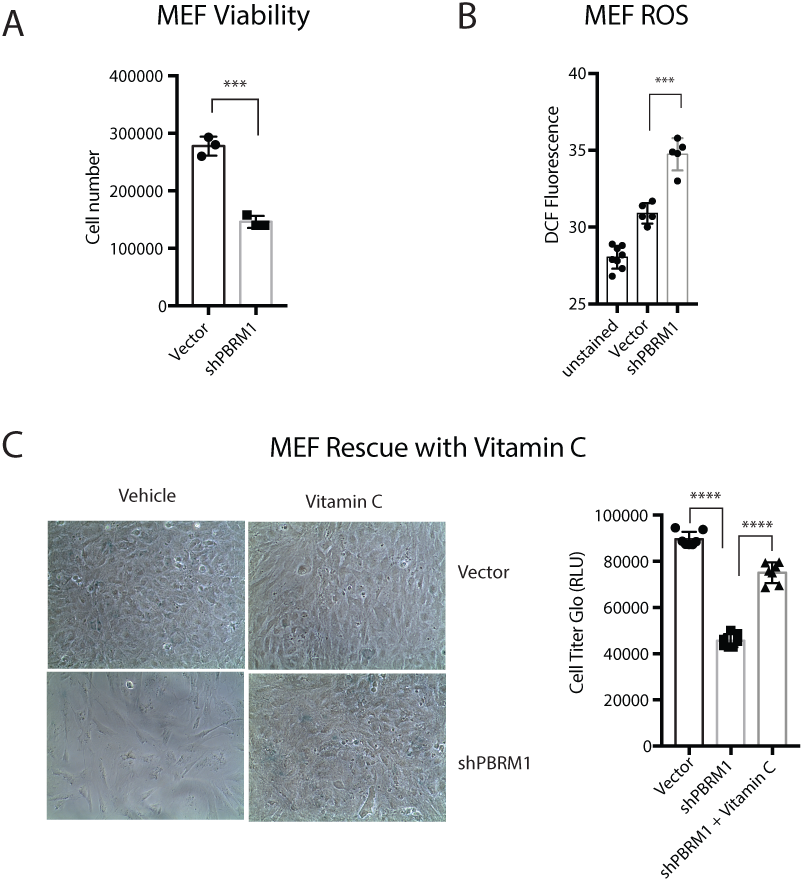
A) Mouse embryonic fibroblasts (MEFs) were counted after 72h growth using trypan blue and data presented as mean ± SD. n = 3. **B**) Equal numbers of MEFs were plated in 96 well format. After 24h the cells were washed with PBS, incubated with H2-DCFDA for 1h and the DCF fluorescence was measured in live cells. Data presented as mean ± SD. n = 5. **C**) MEFs were cultured for 8 days in normal media or media supplemented with 20 *μ*g/mL Vitamin C. Luminescence was measured using CellTiter-Glo^®^ assay system and data presented as mean ± SD. n = 7. A designation of *=p <0.05, ** = p <0.01, *** = p < 0.001, **** = p < 0.0001 (paired Student’s t test). ns, not significant. Error bars represent S.D.

### PBRM1 can play different roles during cancer progression

We next sought to examine how intrinsic changes during oncogenic transformation could alter dependency on PBRM1. To do this we employed the MCF10A series of breast cancer cell lines (Fig 7A). MCF10A cells are non-transformed human mammary epithelial cells (Soule et al., 1990), the MCF10A-NeoT cells are transformed with T24-HRas and passaged in a mouse to make the MCF10A-T1k cells (Dawson et al., 1996). The T1k line produces tumors in about 20% of mice and one of these tumors was passaged in mice to produce the tumorigenic cell line, MCF10A-CA1h, which produces non-metastatic glandular tumors in mice (Santner et al., 2001). We knocked down PBRM1 in each of these cell lines (SI6A) and found dramatically different effects on viability. Similar to other epithelial cell lines, PBRM1 knockdown in MCF10A results in an increase in proliferation and a slight increase in ROS (Fig 7B-left). In contrast, PBRM1 knockdown in the MCF10A-T1k cell line is highly deleterious to viability, causing cells to cease proliferation altogether within 3-4 passages (Fig 7B-right). Similar to MEFs, PBRM1 knockdown in this line induces a highly significant increase in ROS levels, and similar to MEFs, Vitamin C administration can partially restore proliferative capacity in the PBRM1 knockdown (Fig 7C). Upon further progression to malignancy with the CA1h cancer line, shPBRM1 again increases proliferation, acting again as a tumor suppressor (Fig 7D). The differential effects PBRM1 can have on proliferation rates at different stages of transformation was similarly observed for BRG1 in pancreatic cancer (Roy et al., 2015) and ARID1A in hepatocellular carcinoma (Sun et al., 2017). Due to the intrinsic replicative and metabolic stress during a certain stage of transformation, PBRM1 can act as a tumor suppressor or an oncogene. To support this possibility, the MCF10A-T1k cell line, which is the most dependent on PBRM1 for viability also displays an increased level of ROS at baseline compared to MCF10A cells (SI6B). In summary, our data up to this point establish that PBRM1 knockdown can have different effects on viability in the same cell line due to different external stress environments, or in two different cell lines due to cell-type susceptibilities to stress.

**Figure 7:**
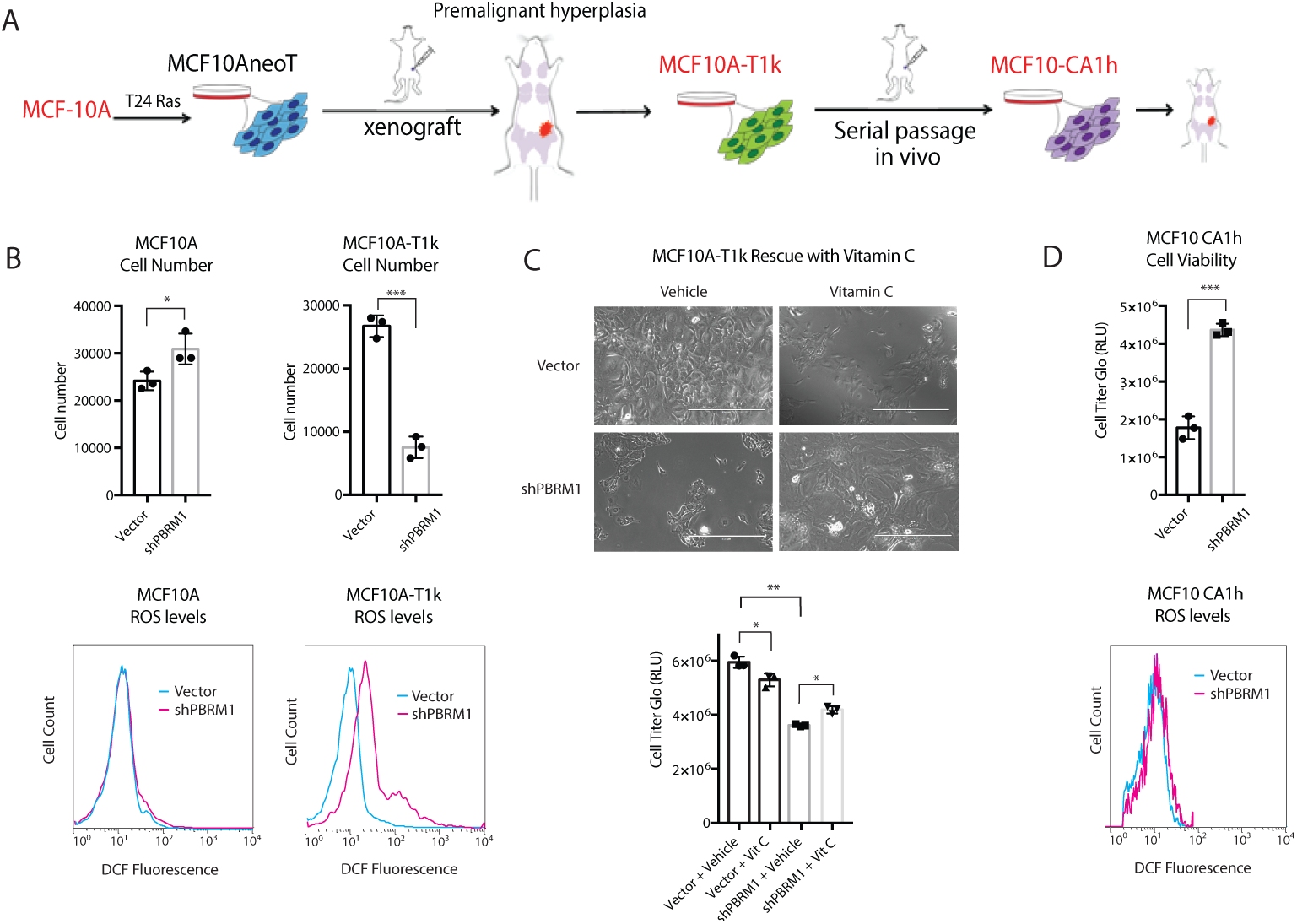
Dual roles for PBRM1 during different stages of tumor progression. **A**) Depiction of MCF10A series of cell lines isolated during different stages of tumor progression. **B**) (top) Human mammary epithelial cell line MCF10A and transformed cell lines MCF10A-T1k were counted after 72h growth using trypan blue and data presented as mean ± SD. n = 3. * p <0.05, *** p<0.001. (bottom) MCF10A and MCF10A-T1k cells were trypsinized and stained for 30 min with H_2_-DCFDA, washed with PBS, and 100,000 cells were analyzed using flow cytometry. **C**) MCF10A-T1k cells were cultured for 7 days in normal media or media supplemented with 20 *μ*g Vitamin C. Luminescence was measured using CellTiter-Glo^®^ assay system and data presented as mean ± SD. n = 3. *p<0.05 ** p<0.01. **D**) (top) Breast cancer cell line MCF10CA1h was cultured for 72h. Luminescence was measured using CellTiter-Glo^®^ assay system and data presented as mean ± SD. n = 3. ** p <0.01, *** p<0.001. (bottom) MCF10CA1h cells were trypsinized and stained for 30 min with H_2_-DCFDA, washed with PBS, and 10,000 cells were analyzed using flow cytometry.

## Discussion

Chromatin regulators are frequently misregulated in cancer with resulting alterations in gene transcription; however, many of these regulators alter a large number of genes to a small degree and can regulate very different sets of genes in different cell lines. An additional challenge resides in the fact that many chromatin regulators modulate transcription differently depending on environmental inputs (Johnson and Dent, 2013). Thus, it has been incredibly challenging to decipher how the transcriptional effects of chromatin regulators observed in a particular cell line relates to its general biochemical function or its phenotype *in vivo*. Traditional cell culture models are often devoid of the environmental stimuli chromatin regulators normally sense, making it a significant challenge to develop a relevant cell culture model for accurately studying these regulators. Here we have used transcriptional analysis of epithelial cells with PBRM1 knockdown to identify pathways involved in epithelial cell maintenance and stress response. In addition, we have validated a role for PBRM1 in the maintenance of epithelial cell identity and identified KLF, AP-1 and ETS transcription factors consensus sequences in both genes downregulated upon shPBRM1, as well as regions with decreased chromatin accessibility upon shPBRM1. From this, we have validated a role for PBRM1 in restraining ROS production and inducing both apoptotic and cell survival pathways under high stress conditions. (Roupé et al., 2014; Y. Wang et al., 2014).

In addition to changes in transcription factor expression and localization during cellular stress, oxidative stress and metabolic stress are known to specifically upregulate H3K14Ac at stress response genes (Schram et al., 2013), a histone mark specifically recognized by PBRM1 (Porter and Dykhuizen, 2017). While H3K14Ac is generally found at active promoters with H3K9Ac, it is found without H3K9Ac at inducible genes (Karmodiya et al., 2012) and is specifically increased in gene bodies during stress (Johnsson et al., 2009). H3K14Ac is also a mark associated with renal epithelium adaption to oxidative stress (Mahalingaiah et al., 2016; 2015), high-fat diet induced inflammation in rodents (Suter et al., 2012), endoplasmic reticulum stress (Dicks et al., 2015), and sites of DNA-damage (Chiu et al., 2017), a process for which PBRM1 has a well-established role (Kakarougkas et al., 2014). Therefore, specific patterns of histone acetylation likely delineate a subset of stress-response genes targeted by PBRM1, which is likely to be unique for a particular cell type, as well as a particular stressor.

A role for PBRM1 in stress response is in agreement with recent findings that PBRM1 deletion alone is not sufficient for transformation but acts to facilitate oncogenesis in cooperation with VHL deletion (Espana-Agusti et al., 2017; Gu et al., 2017; Nargund et al., 2017). PBRM1 deletion allows for an amplification of oncogenic signaling (Gao et al., 2017; Nargund et al., 2017), as well as a bypass of checkpoints induced by replication stress after VHL deletion (Espana-Agusti et al., 2017). This could be in part due to PBRM1’s role in regulating the hypoxia stress response in cooperation with HIF1a, which allows for the reduction of ROS and induction of apoptosis in response to hypoxia (Kim et al., 2006).

Not only is a role in stress response likely part of PBRM1’s function as a tumor suppressor during cancer initiation, it may also be involved in PBRM1’s protective function against cancer therapeutics, similar to the protective role PBRM1 plays under high stress conditions. ccRCC patients with PBRM1 mutations tend to have favorable prognosis (Piva et al., 2015) and recent studies indicate that PBRM1 mutant tumors respond particularly well to sunitinib (Beuselinck et al., 2017) and PD-1 inhibitors (Miao et al., 2018). In support of this, a recent CRISPR-Cas9 screen identified PBRM1, along with other PBAF-specific subunits, as resistance factors against T-cell mediated killing (Pan et al., 2018). This is related to a general role for PBRM1 in suppressing the inflammatory response, as PBRM1 deletion also increases innate immunity hyperinflammation in the gut (He et al., 2017; Shu et al., 2017). The general role for PBRM1 in the stress response could relate to PBRM1’s role in suppressing inflammation (and T-cell mediated toxicity) through the regulation of homeostasis (Chovatiya and Medzhitov, 2014), although that connection will need to be explored further.

## Author Contributions

EGP, AD, BC, and ECD conceived the experiments. EGP, AD, BC, MKW, and ECD designed experiments. EGP, AD, BC, JCS and HL performed experiments. BC and BCC performed bioinformatics analyses. MK provided advice for ATAC-Seq analysis. EGP, AD, BC, BCC, and ECD wrote the manuscript.

## Acknowledgements

This research was supported with grants from the NIH (U01CA207532 for ECD, R01CA207751 for MWK, K22HL125593 for MK). This work was supported by the Office of the Assistant Secretary of Defense for Health Affairs, through the Peer Reviewed Cancer Research Program, under Award No. W81XWH-17-1-0267 to ECD. Opinions, interpretations, conclusions and recommendations are those of the author and are not necessarily endorsed by the Department of Defense. ECD was supported by The V Foundation for Cancer Research (V2014-004 and D2016-030). EGP was supported by the Borch Graduate Fellowship from the Purdue Department of Medicinal Chemistry and Molecular Pharmacology. Additional support was from the Purdue University Center for Cancer Research, NIH grant P30 CA023168.

## Materials and Methods

### Cell culture

HK-2 were cultured in RPMI (Corning Mediatech) supplemented with 10% fetal bovine serum (J R Scientific), 1% antibiotics (100 units/ml penicillin and 100 g/ml streptomycin; Corning Mediatech), and 1% L -glutamine (Corning Mediatech) at 37 °C in a humidified atmosphere in a 5% CO2 incubator.

MEF cells were cultured in DMEM supplemented with 10% fetal bovine serum (J R Scientific), 1% antibiotics (100 units/ml penicillin and 100g/ml streptomycin; Corning Mediatech), 1% nonessential amino acids (Corning Mediatech), 1% L -glutamine (Corning Mediatech) and 0.1% β-mercaptoethanol (Gibco, Thermo Scientific) at 37 °C in a humidified atmosphere in a 5% CO2 incubator.

MCF10A cells were cultured in 1:1 DMEM (Corning Mediatech, Inc.) and F12 (Corning Mediatech, Inc.) supplemented with 29 mM Hepes (Amresco, LLC), 10 mM Sodium Bicarbonate (Macron), 5% Horse serum (Sigma), 10 *μ*M/mL Insulin (Sigma), 10ng/mL Epidermal Growth Factor (EGF) (Gold Biotechnology), 0.5 *μ*g/mL hydrocortisone (Sigma), 100 ng/mL cholera toxin (Sigma), and 1% antibiotics (100 units/ml penicillin and 100 g/ml streptomycin; Corning Mediatech) at 37 °C in a humidified atmosphere in a 5% CO_2_ incubator.

MCF10A T1K, MCF10A CA1h, were cultured in DMEM (Corning Mediatech) supplemented with 10% fetal bovine serum (J R Scientific) 1% L -glutamine (Corning Mediatech), 1% antibiotics (100 units/ml penicillin and 100 g/ml streptomycin; Corning Mediatech), and 1% Sodium Pyruvate (Corning Mediatech) at 37 °C in a humidified atmosphere in a 5% CO_2_ incubator.

NMuMG cells were grown in DMEM (Corning Mediatech) supplemented with 10% fetal bovine serum (J R Scientific), 10 *μ*M/mL Insulin (Sigma), 1% L -glutamine (Corning Mediatech), 1% antibiotics (100 units/ml penicillin and 100 g/ml streptomycin; Corning Mediatech), and 1% Sodium Pyruvate (Corning Mediatech) at 37 °C in a humidified atmosphere in a 5% CO_2_ incubator.

MDCK cells were cultured in in DMEM (Corning Mediatech) supplemented with 10% fetal bovine serum (J R Scientific), 1% L -glutamine (Corning Mediatech), 1% antibiotics (100 units/ml penicillin and 100 g/ml streptomycin; Corning Mediatech), 1% nonessential amino acids (Corning Mediatech), 10 mM Hepes (HyClone) and 1% Sodium Pyruvate (Corning Mediatech), at 37 °C in a humidified atmosphere in a 5% CO_2_ incubator.

All the media were supplemented with 1:10,000 dilution of Plasmocin™ (InvivoGen).

### Cell culture and treatments

Cells were seeded 24-72 h before treatment such that they were 50-80 % confluent at the time of experiment. For hydrogen peroxide treatment, indicated concentrations of freshly prepared hydrogen peroxide were added to the treatment groups for the indicated time periods in their regular media. For glucose starvation studies, the regular media was replaced with glucose free media for the indicated time periods. Following the completion of treatment, cells were washed once with PBS, harvested by trypsinization and either processed immediately or flash frozen and stored at -80 °C for future use.

### Generation of cell lines

Knockdown was performed using shRNA-mediated knockdown with lentiviral construct pLKO.1. The shRNA constructs contain the following mature antisense sequences: Human PBRM1: (TRCN0000015994, TTTGTAGATCAAAGACTCCGG Mouse PBRM1 (TRCN0000081820) TTCTAGGTTGTATGCCTGTCG

Mouse Brd7 Clone ID: TRCN0000030015 ATAATCATGGAGTAGCCAGGC

Mouse Brg1 TRCN0000071386 TTCTCAATAATGTGTCGGGCG

Mouse Arid1a (TRCN0000071395, Origene TG517733), ATTGTAGGTCATGTCATTTCG Canine PBRM1-1: ACATCATCATACTCTTCCA

Canine PBRM1-2: ACCAACAGCCATACAACCA

PBRM1-TetO-Fuw and Vector-TetO-Fuw were used with pLenti CMV rtTA3 Hygro (w785-1) (a gift from Eric Campeau, Addgene plasmid number 26730) for tetracycline inducible expression.

*Lentiviral Infection*-HEK293T cells were transfected with lentivirus constructs along with packaging vectors pMD2.G and psPAX2. After 48 h, the supernatant was collected and concentrated by ultracentrifugation (17,300 rpm for 2 h) and resuspended in 200 *μ*l of PBS. Cells were infected with concentrated virus using spinfection (1500 rpm in swing bucket centrifuge for 1 h). Fresh medium was added 16 h after infection, and cells were allowed to recover for 24 h before selection. Cells were selected for 2 weeks with puromycin (0.6 *μ*g/ml) (Sigma-Aldrich) and hygromycin (200 *μ*g/ml) (Corning Mediatech). Cells were cultured with 2 *μ*g/ml doxycycline (EMDChemicals, San Diego, CA) for 72 h prior to experiments to induce protein expression.

### 3D culture

Cells were embedded between 2 layers of Cultrex^®^ Basement Membrane Extract (BME) (R&D Systems) on 8-well Chamber Slide. Wells were pre-coated with BME (200 *μ*l/well) to allow polymerization at 37°C for 15 minutes. Cells were then seeded at 20,000 cells/well density. After attachment (30 minutes at 37°C), cells were covered with a second layer of BME/culture medium (1:19, 5%) to polymerize overnight at 37°C. Cells were incubated for 10 days, and the medium was replenished every 3 days. At the end of incubation, cells were fixed and subjected to immunofluorescence analysis.

### Immunofluorescence staining

Cells were washed twice with ice-cold PBS, added 2-3 volumes of ice-cold PBS-EDTA and shaken on ice for 15-30 minutes. BME was detached from the bottom of culture surface by gently scraping the bottom with a pipette tip. The solution was transferred to a conical tube and gently shaken on ice for 15-30 minutes. When BME was dissolved completely, the solution was centrifuged at 120g for 1-2 minutes. The supernatant was carefully aspirated and cells were gently resuspended in the remaining supernatant. Pipetted approximated 15 *μ*l of the cells suspension onto a glass bottom dish, allowed cells to settle and adhere to the glass. Cells were fixed using formalin for 20 minutes at room temperature (RT). Next, cells were permeabilized with 0.5% Triton X-100 in PBS for 5 minutes at RT and washed 3 times with 100 mM glycine in PBS at RT. Fixed cells were blocked for 1.5 hours with 10% goat serum. Cells were incubated overnight at 4°C with primary antibodies. The primary antibodies used were as follows: rat anti– α_6_-integrin (Millipore; 1:100 in 0.2% Triton X-100, 0.1% BSA, 0.05% Tween 20 in PBS) and rabbit anti–Zo-1 (Invitrogen, 1:100 in 0.2% Triton X-100, 0.1% BSA, 0.05% Tween 20 in PBS). Cells were incubated with secondary antibody for 1 hour, followed by 3 washes at RT. Secondary antibodies were as follows: FITC goat anti-rat and Biotin-SP-conjugated AffiniPure goat anti-rabbit (Jackson ImmunoResearch). Cells were incubated with Texas Red Avidin D (Vector) for 1hour. Cell nuclei were counterstained with DAPI for 10 minutes and washed 3 times with PBS. Cells were incubated in PBS and imaged by confocal microscopy.

### Confocal microscopy

Confocal laser scanning microscopy experiments were conducted using the Zeiss LSM 880 Upright Confocal.

### Top Flash Reporter Assay

NMuMG cells were transfected with 10:1 ratio of M50 Super 8x Top flash (Addgene 12456) to pcDNA3.1.CMV-renilla-Neo. The cells were transfected using Lipofectamine with 3:1 ratio of total DNA to lipofectamine reagent. After 24 h, the cells were trypsinized and 20,000 cells/well were plated in white 96-well tissue culture treated plates. After an additional 24 h of growth, the firefly and renilla luciferase levels were measured using the Dual Glo^®^ assay system (Promega).

### Annevin V Apoptosis detection in NMuMG cells

NMuMG PLKO and NMuMG shPBRM1 cells were seeded at a density of ~ 1.5 × 10^6^ cells/60mm dish and cultured for 24 h. The cells were then given treatments of 200 *μ*M H_2_O_2_ in media for 0-4 h, followed by cell harvesting using Accutase (Innovative cell technologies, CA, USA) and apoptosis detection using the FITC Annexin V apoptosis detection kit (BD Pharmingen, Cat. # 556547) as per the manufacturer’s instructions. The cells were immediately analyzed by flow cytometry using the Guava EasyCyte Benchtop Flow Cytometer (Millipore Sigma, MA, USA). The results were analyzed using FlowJo software.

### Immunoblotting

NMuMG PLKO and NMuMG shPBRM1 cells were seeded in 60 mm dishes at a density of ~ 1.5 × 10^6^ cells/dish, Caki2+Vector and Caki2+PBRM1 cells were seeded in 6-well plates at a density of ~ 6 × 10^4^ cells/well, and given H_2_O_2_ treatments as described before for the indicated time periods, followed by cell harvesting by trypsinization. Whole cell extracts were prepared by dissolving the cell pellets in RIPA buffer (50 mM Tris (pH 8.0), 150 mM NaCl, 0.1% SDS, 0.5% Na Deoxycholate, 1% NP-40) supplemented with freshly added PMSF, aprotinin, leupeptin and pepstatin, and incubation for 30 min at 4 °C. The lysates were centrifuged at 13000 × g for 30 min at 4 °C and the supernatants were preserved. Protein concentration estimations for the supernatants were done using BCA protein assay kit (Pierce Biotechnology, IL, USA) using BSA as standard and 40 *μ*g of whole cell extracts were run on a 4-12 % bis-tris gradient protein gel, transferred to PVDF membrane and probed with primary antibodies in 5% BSA at 4 °C for 16 h.

### Antibodies

Cleaved PARP (Asp214) (7C9) (Mouse specific) Cell Signaling Technology-#9548, PBRM1 (Bethyl Laboratories-PBRM1 Antibody, #A301-591A)

β-actin (Santa Cruz Biotechnology, #sc-47778)

BRG1 (G-7) Santa Cruz sc-17796

BRD7 Bethyl A302-304A

Vimentin BD Biosciences 550513

E Cadherin BD Biosciences 610182 GAPDH (6C5) (Santa Cruz sc-32233)

Cleaved PARP (Asp214) (Human Specific) Cell Signaling #9541

LaminB (A-11) sc-377000

### H_2_-DCFDA staining for intracellular ROS using flow cytometry

NMuMG PLKO and NMuMG shPBRM1 cells were seeded at a density of ~ 2 × 10^6^ cells/60mm dish. The cells were cultured for 48 h, harvested by trypsinization, washed once with PBS and stained for intracellular ROS by incubation with freshly prepared 10 *μ*M H2-DCFDA (Invitrogen, Cat. # D399) in PBS for 30 min at 37 °C in dark. Following the incubation, the cells were centrifuged at 250 x g for 5 min, resuspended in PBS and immediately analyzed by flow cytometry. The results were analyzed using FlowJo software.

### Stress treatments and H_2_-DCFDA staining using microplate reader

NMuMG cells were seeded at a density of 6.0 × 10^4^ in a 96-well tissue culture plate. The cells were cultured for 24 h, following which they were subjected to the following stress conditions: H_2_O_2_ treatment (0-800 *μ*M) for 2h followed by 10 min recovery in PBS, glucose starvation by culturing in various glucose concentrations for 16h, CoCl2 treatment (0-500*μ*M) for 24h or doxorubicin treatment (0-10 *μ*M) for 24h. At the end of the treatments, cells were washed once with PBS, and stained with freshly prepared 10 *μ*M H_2_-DCFDA (Invitrogen, Cat. # D399) in PBS for 30 min at 37 °C in dark. Cells were washed again 2x with PBS and fluorescence measurements were taken using a microplate reader at excitation/emission wavelengths of 485/530 nm. Unstained cells were used as the negative controls.

### H_2_O_2_ detection assay

H_2_O_2_ levels were measured using Amplex^®^ Red Hydrogen Peroxide/Peroxidase Assay kit (Invitrogen). The H_2_O_2_ levels were determined from whole cell lysates (in RIPA) according to manufacturer’s instruction. The concentration of H_2_O_2_ was determined for lysates generated from 5,000 and 10,000 cells in 50 *μ*L by plotting fluorescence levels against experimentally determined dose curves.

### Viability assays using CellTiter-Glo^®^

Cells were plated in 96-well white tissue culture plates and cultured for the indicated time. Antioxidant rescue experiments were performed with fresh media daily containing 20 *μ*g/mL Vitamin C or 250 *μ*M N-acetylcysteine (NAC). CellTiter-Glo assay reagent was added directly to cells, incubated for 20 min, and the luminescence was measured on a GloMax^®^ microplate reader.

### LDH assays using LDH Cytotoxicity Assay Kit II from Abcam (ab65393)

NMuMG cells were seeded at a density of 6.0 × 10^4^ in 96 well tissue culture plates. The cells were cultured for 24 h, following which they were subjected to H_2_O_2_ treatment (0-800 *μ*M) for 6h or 24h. Media was harvested from wells (10 *μ*L) and transferred to a separate 96 well assay plate along with negative control (media alone) and positive control (lysed cells). LDH Reaction Mix (100 *μ*l) was added to each well, mixed and incubated for 30 min at room temperature. 7. The absorbance at 450 nm was measured on the GloMax^®^ microplate reader.

### H_2_-DCFDA staining for MCF10A series followed by flow cytometry

MCF10A, MCF10A-T1K, MCF10A-CA1h and MCF10A-CA1a PLKO and their respective PBRM1 knockdown cells were seeded in 60 mm dishes in MCF10A media and cultured for 48 h such that they reach 50-80 % confluency at the day of the experiment. The cells were then harvested using trypsin, washed once using serum-free and phenol red-free media and stained for intracellular ROS with freshly prepared 10 *μ*M H_2_-DCFDA in PBS –Glucose (1X PBS supplemented with 25 mM glucose) as described before. The cells were immediately examined by flow cytometry and the results were analyzed using FlowJo software.

### RNA-seq

RNA isolation, library construction, sequencing and transcriptome analysis was performed as described in our previous publication (Chowdhury et al. 2016). Sequencing was performed in biological triplicate except for Caki2 in 3D culture, which was performed in biological duplicate. RNA-seq of Caki1 and Caki2 grown in 3D culture was performed at Beijing Genomics Institute (BGI) using Illumina HiSeq technology, while RNA-Seq of the HK2 and NMuMG epithelial cell lines was performed at the Purdue Genomics Core using Illumina HiSeq technology. The resulting reads were trimmed using Trimmomatic utility (Bolger et al., 2014) and mapped to hg19 or mm9 using STAR (Dobin et al., 2013) using default parameters. Read counts were obtained using HTSeq-count (Anders et al., 2015) in conjunction with a standard gene annotation files from UCSC (University of California Santa Cruz; http://genome.ucsc.edu) and differential expression was determined using DESeq2 pipeline (Love et al., 2014). Differentially expressed genes were filtered using a false discovery rate threshold of < 0.05 and a fold change threshold of > 1.3-fold relative to the reference sample. Gene ontology and transcription factor prediction analyses were performed using GeneCodis (Nogales-Cadenas et al., 2009). Data sets generated in these experiments are available at the Gene Expression Omnibus under accession number GSE113606.

### ATAC-seq

The ATAC-seq protocol originally described (Buenrostro et al., 2015) was adapted as follows for HK2 and NMuMG isogenic lines: 150,000 cells were resuspended in Nuclei Lysis buffer containing 0.05% IGEPAL CA-630, incubated for 5 minutes on ice and centrifuged for 10 minutes at 500xg at 4 °C. Nuclei extraction was confirmed by microscopic inspection and the nuclei pellet was resuspended in transposition master mix. Tagmentation, cleanup of tagmented DNA, and PCR enrichment was performed as per original description. High throughput sequencing was performed by HiSeq2500 using 50 bp paired-end at the Purdue Genomics Core. Sequenced reads were mapped by the Bowtie2 aligner (Ben Langmead et al., 2009) using hg19 or mm10 reference genome respectively. Reads mapping to the mitochondrial genome were discarded. Bigwig files were generated for visual inspection of tracks using the bamCoverage utility of deepTools (Ramírez et al., 2016). Peaks of differential accessibility were identified using the SICER-df-rb utility (Xu et al., 2014) with a false discovery rate threshold of < 0.05 and a fold change threshold of > 1.5-fold difference in accessibility. Scaled heat maps were generated for the peak regions using the computeMatrix and plotHeatmap utilities of deepTools. Peak regions were analyzed for enrichment of sequence motifs and association with genomic elements using the findMotifs and annotatePeaks utilities of HOMER (Heinz et al., 2010). Data sets generated in these experiments are available at the Gene Expression Omnibus under accession number GSE113606.

### qRT-PCR Quantitative PCR

RNA was isolated from cells with Trizol (Ambion, Thermofisher). Total RNA was converted to cDNA with Verso cDNA Sythesis Kit (Thermo Scientific). Real-time PCR was performed using a Bio-Rad CFX Connect Real-Time system and Thermo Scientific Maxima SYBR Green qPCR Master Mix (Thermo Scientific). The results were analyzed using the Pfaffl method.(Pfaffl, 2001)

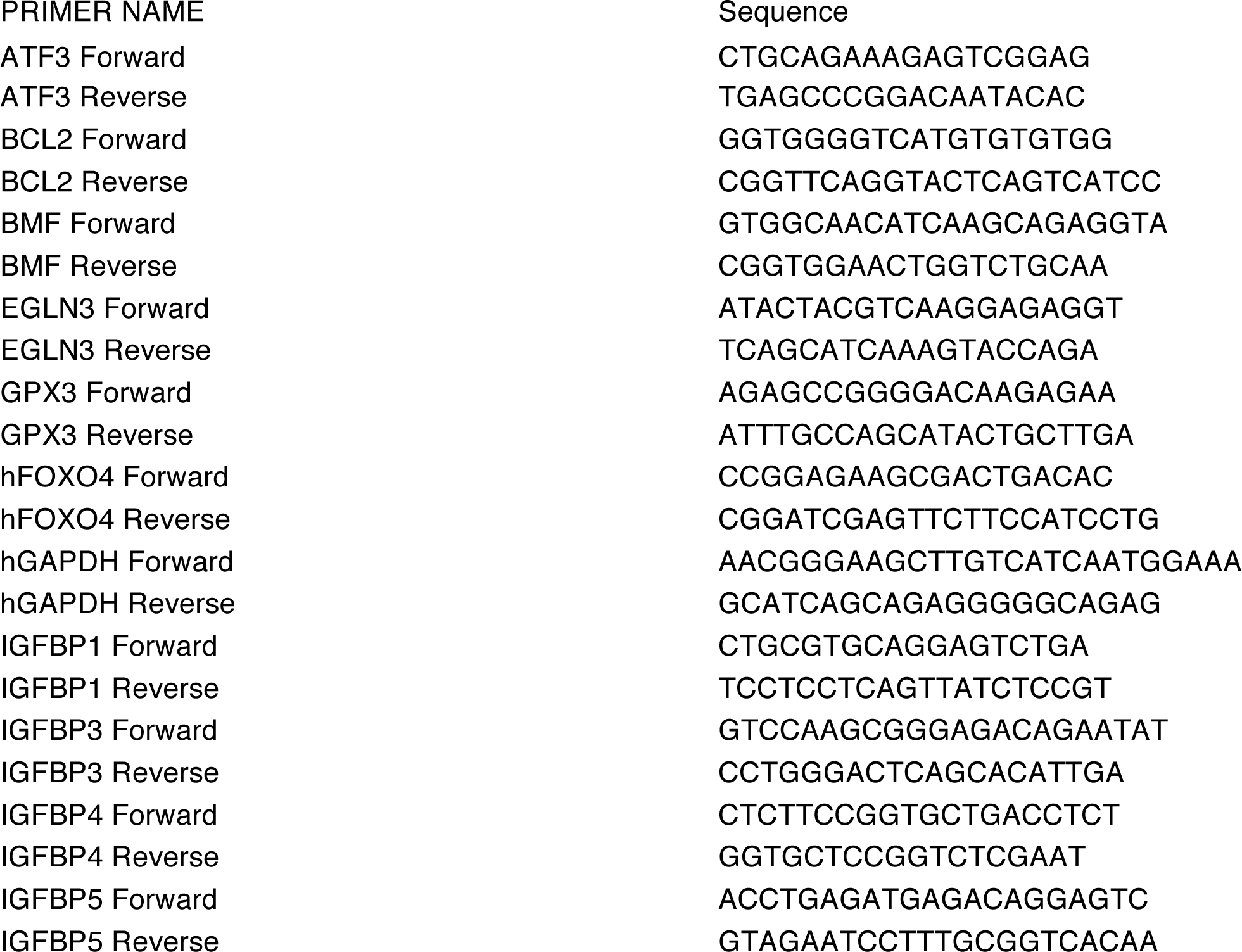
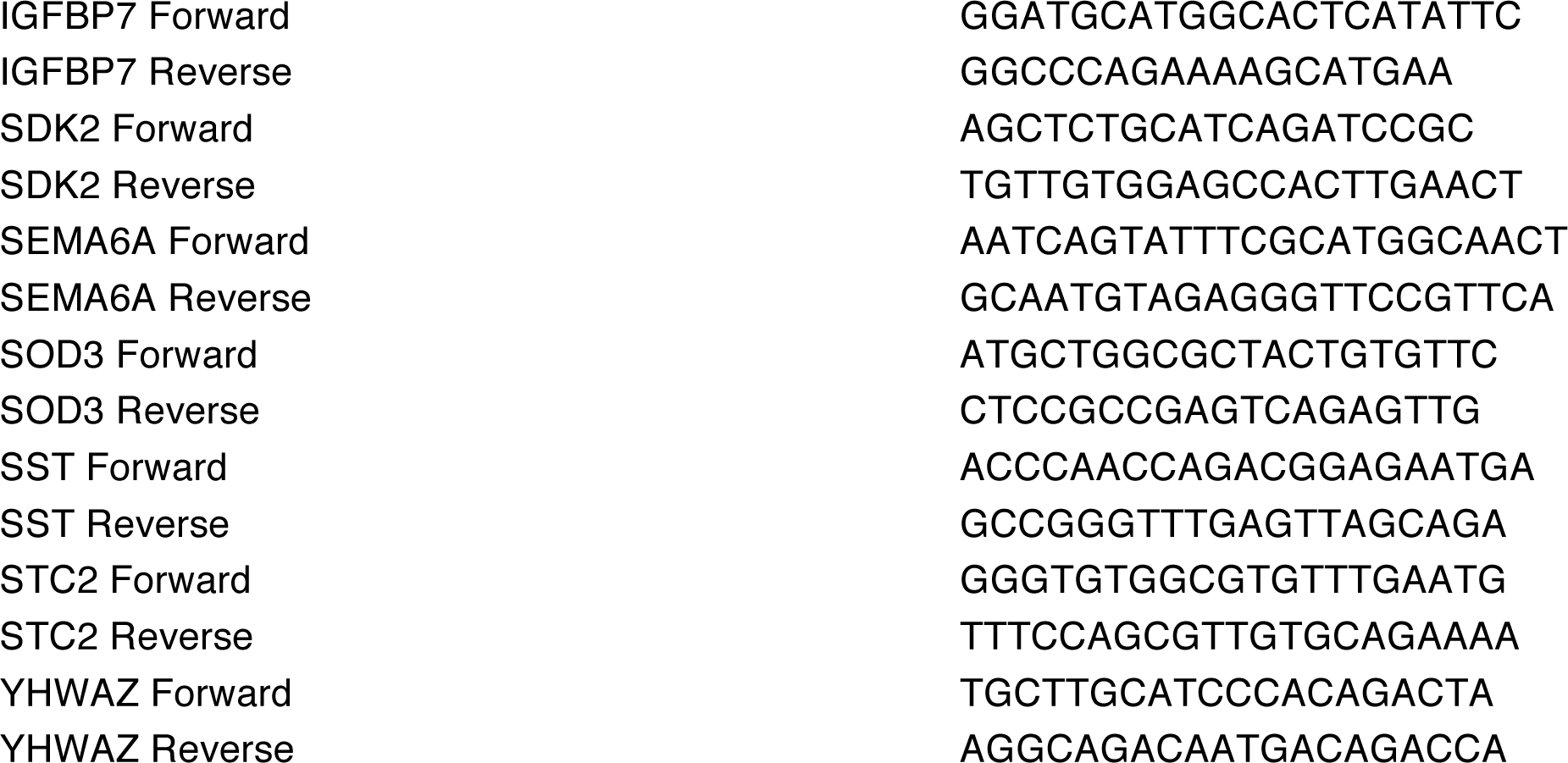

## References

Anders, S., Pyl, P.T., Huber, W., 2015. HTSeq--a Python framework to work with high-throughput sequencing data. Bioinformatics 31, 166–169. doi:10.1093/bioinformatics/btu638

Beagan, J.A., Duong, M.T., Titus, K.R., Zhou, L., Cao, Z., Ma, J., Lachanski, C.V., Gillis, D.R., Phillips-Cremins, J.E., 2017. YY1 and CTCF orchestrate a 3D chromatin looping switch during early neural lineage commitment. Genome Research 27, 1139–1152. doi:10.1101/gr.215160.116

Ben Langmead, Trapnell C., Pop, M., Salzberg, S.L., 2009. Ultrafast and memory-efficient alignment of short DNA sequences to the human genome. Genome Biol 10, R25. doi:10.1186/gb-2009-10-3-r25

Berglund, F.M., Weerasinghe, N.R., Davidson, L., Lim, J.C., Eickholt, B.J., Leslie, N.R., 2012. Disruption of epithelial architecture caused by loss of PTEN or by oncogenic mutant p110α/PIK3CA but not by HER2 or mutant AKT1. Oncogene 32, 4417–4426. doi:10.1038/onc.2012.459

Beuselinck, B., Verbiest, A., Couchy, G., Job, S., de Reynies, A., Meiller, C., Albersen, M., Verkarre, V., Lerut, E., Méjean, A., Patard, J.-J., Laguerre, B., Rioux-Leclercq, N., Schöffski, P., Oudard, S., Zucman-Rossi, J., 2017. Pro-angiogenic gene expression is associated with better outcome on sunitinib in metastatic clear-cell renal cell carcinoma. Acta Oncologica 1–11. doi:10.1080/0284186X.2017.1388927

Bolger, A.M., Lohse, M., Usadel, B., 2014. Trimmomatic: a flexible trimmer for Illumina sequence data. Bioinformatics 30, 2114–2120.

Brownlee, P.M., Chambers, A.L., Cloney, R., Bianchi, A., Downs, J.A., 2014. BAF180 promotes cohesion and prevents genome instability and aneuploidy. CellReports 6, 973–981. doi:10.1016/j.celrep.2014.02.012

Buenrostro, J., Wu, B., Chang, H., Greenleaf, W., 2015. ATAC-seq: A Method for Assaying Chromatin Accessibility Genome-Wide. Current protocols in molecular biology / edited by Frederick M. Ausubel … [et al.] 109, 21.29.1–21.29.9. doi:10.1002/0471142727.mb2129s109

Cancer Genome Atlas Research Network, 2013. Comprehensive molecular characterization of clear cell renal cell carcinoma. Nature 499, 43–49. doi:10.1038/nature12222

Charlop-Powers, Z., Zeng, L., Zhang, Q., Zhou, M.-M., 2010. Structural insights into selective histone H3 recognition by the human Polybromo bromodomain 2. Cell Res 20, 529–538.

Chiu, L.-Y., Gong, F., Miller, K.M., 2017. Bromodomain proteins: repairing DNA damage within chromatin. Philos. Trans. R. Soc. Lond., B, Biol. Sci. 372, 20160286. doi:10.1098/rstb.2016.0286

Chovatiya, R., Medzhitov, R., 2014. Stress, Inflammation, and Defense of Homeostasis. Mol Cell 54, 281–288. doi:10.1016/j.molcel.2014.03.030

Chowdhury, B., Porter, E.G., Stewart, J.C., Ferreira, C.R., Schipma, M.J., Dykhuizen, E.C., 2016. PBRM1 Regulates the Expression of Genes Involved in Metabolism and Cell Adhesion in Renal Clear Cell Carcinoma. PLoS ONE 11, e0153718. doi:10.1371/journal.pone.0153718

Cohet, N., Stewart, K.M., Mudhasani, R., Asirvatham, A.J., Mallappa, C., Imbalzano, K.M., Weaver, V.M., Imbalzano, A.N., Nickerson, J.A., 2010. SWI/SNF chromatin remodeling enzyme ATPases promote cell proliferation in normal mammary epithelial cells. J. Cell. Physiol. 260, /a–n/a. doi:10.1002/jcp.22072

Dawson, P.J., Wolman, S.R., Tait, L., Heppner, G.H., Miller, F.R., 1996. MCF10AT: a model for the evolution of cancer from proliferative breast disease. The American Journal of Pathology 148, 313–319.

de Nadal, E., Ammerer, G., Posas, F., 2011. Controlling gene expression in response to stress. Nature Reviews Genetics 12, 833–845. doi:10.1038/nrg3055

Dicks, N., Gutierrez, K., Michalak, M., Bordignon, V., Agellon, L.B., 2015. Endoplasmic Reticulum Stress, Genome Damage, and Cancer. Front. Oncol. 5, 61. doi:10.3389/fonc.2015.00011

Dobin, A., Davis, C.A., Schlesinger, F., Drenkow, J., Zaleski, C., Jha, S., Batut, P., Chaisson, M., Gingeras, T.R., 2013. STAR: ultrafast universal RNA-seq aligner. Bioinformatics 29, 15–21.

Duan, M.R., Smerdon, M.J., 2014. Histone H3 Lysine 14 (H3K14) Acetylation Facilitates DNA Repair in a Positioned Nucleosome by Stabilizing the Binding of the Chromatin Remodeler RSC (Remodels Structure of Chromatin). J Biol Chem 289, 8353–8363. doi:10.1074/jbc.M113.540732

Espana-Agusti, J., Warren, A., Chew, S.K., Adams, D.J., Matakidou, A., 2017. Loss of PBRM1 rescues VHL dependent replication stress to promote renal carcinogenesis. Nat Commun 8, 2026. doi:10.1038/s41467-017-02245-1

Fulda, S., Gorman, A.M., Hori, O., Samali, A., 2010. Cellular Stress Responses: Cell Survival and Cell Death. International Journal of Cell Biology 2010, 1–23. doi:10.1155/2010/214074

Gao, W., Li, W., Xiao, T., Liu, X.S., Kaelin, W.G., 2017. Inactivation of the PBRM1 tumor suppressor gene amplifies the HIF-response in VHL-/- clear cell renal carcinoma. P Natl Acad Sci Usa 114, 1027–1032. doi:10.1073/pnas.1619726114

Geou-Yarh Liou, P.S., 2010. Reactive oxygen species in cancer. Free radical research 44, 479–496. doi:10.3109/10715761003667554

Gerlinger, M., Horswell, S., Larkin, J., Rowan, A.J., Salm, M.P., Varela, I., Fisher, R., McGranahan, N., Matthews, N., Santos, C.R., Martinez, P., Phillimore, B., Begum, S., Rabinowitz, A., Spencer-Dene, B., Gulati, S., Bates, P.A., Stamp, G., Pickering, L., Gore, M., Nicol, D.L., Hazell, S., Futreal, P.A., Stewart, A., Swanton, C., 2014. Genomic architecture and evolution of clear cell renal cell carcinomas defined by multiregion sequencing. Nat Genet 46, 225–233. doi:10.1038/ng.2891

Ghaleb, A.M., McConnell, B.B., Kaestner, K.H., Yang, V.W., 2011. Altered intestinal epithelial homeostasis in mice with intestine-specific deletion of the Krüppel-like factor 4 gene. Developmental Biology 349, 310–320. doi:10.1016/j.ydbio.2010.11.001

Gu, Y.-F., Cohn, S., Christie, A., McKenzie, T., Wolff, N.C., Do, Q.N., Madhuranthakam, A., Pedrosa, I., Wang, T., Dey, A., Busslinger, M., Xie, X.-J., Hammer, R.E., McKay, R.M., Kapur, P., Brugarolas, J., 2017. Modeling Renal Cell Carcinoma in mice: Bap1 and Pbrm1 Inactivation Drive Tumor Grade. Cancer Discovery CD–17–0292.

Hall, H.G., Farson, D.A., Bissell, M.J., 1982. Lumen formation by epithelial cell lines in response to collagen overlay: a morphogenetic model in culture. P Natl Acad Sci Usa 79, 4672–4676.

He, X., Yu, J., Wang, M., Cheng, Y., Han, Y., Yang, S., Shi, G., Sun, L., Fang, Y., Gong, S.-T., Wang, Z., Fu, Y.-X., Pan, L., Tang, H., 2017. Bap180/Baf180 is required to maintain homeostasis of intestinal innate immune response in Drosophila and mice. Nature Microbiology 2017 2:6 2, 17056. doi:10.1038/nmicrobiol.2017.56

Heinz, S., Benner, C., Spann, N., Bertolino, E., Lin, Y.C., Laslo, P., Cheng, J.X., Murre, C., Singh, H., Glass, C.K., 2010. Simple combinations of lineage-determining transcription factors prime cis-regulatory elements required for macrophage and B cell identities. Mol Cell 38, 576–589. doi:10.1016/j.molcel.2010.05.004

Huang, L., Peng, Y., Zhong, G., Xie, W., Dong, W., Wang, B., Chen, X., Gu, P., He, W., Wu, S., Lin, T., Huang, J., n.d. PBRM1 suppresses bladder cancer by cyclin B1 induced cell cycle arrest. Oncotarget 5.

Huang, X., Gao, X., Diaz-Trelles, R., Ruiz-Lozano, P., Wang, Z., 2008. Coronary development is regulated by ATP-dependent SWI/SNF chromatin remodeling component BAF180. Developmental Biology 319, 258–266. doi:10.1016/j.ydbio.2008.04.020

Lee, H., Dai, F., Zhuang, L., Xiao, Z.-D., Kim, J., Zhang, Y., Ma, L., You, M.J., Wang, Z., Gan, B., 2016. BAF180 regulates cellular senescence and hematopoietic stem cell homeostasis through p21. Oncotarget 5. doi:10.18632/oncotarget.8102

Jiang, S., Zhang, E., Zhang, R., Li, X., 2016. Altered activity patterns of transcription factors induced by endoplasmic reticulum stress. BMC Biochemistry 17, 33. doi:10.1186/s12858-016-0060-2

Johnson, D.G., Dent, S.Y.R., 2013. Chromatin: Receiver and Quarterback for Cellular Signals. Cell. doi:10.1016/j.cell.2013.01.017

Johnsson, A., Durand-Dubief, M., Xue-Franzén, Y., Rönnerblad, M., Ekwall, K., Wright, A., 2009. HAT-HDAC interplay modulates global histone H3K14 acetylation in gene-coding regions during stress. EMBO Rep 10, 1009–1014. doi:10.1038/embor.2009.127

Kadoch, C., Hargreaves, D.C., Hodges, C., Elias, L., Ho, L., Ranish, J., Crabtree, G.R., 2013. Proteomic and bioinformatic analysis of mammalian SWI/SNF complexes identifies extensive roles in human malignancy. Nat Genet 45, 592–601. doi:10.1038/ng.2628

Kaeser, M.D., Aslanian, A., Dong, M.-Q., Yates, J.R., Emerson, B.M., 2008. BRD7, a novel PBAF-specific SWI/SNF subunit, is required for target gene activation and repression in embryonic stem cells. J Biol Chem 283, 32254–32263. doi:10.1074/jbc.M806061200

Kakarougkas, A., Ismail, A., Chambers, A.L., Riballo, E., Herbert, A.D., Künzel, J., Löbrich, M., Jeggo, P.A., Downs, J.A., 2014. Requirement for PBAF in Transcriptional Repression and Repair at DNA Breaks in Actively Transcribed Regions of Chromatin. Mol Cell 55, 723–732. doi:10.1016/j.molcel.2014.06.028

Kalluri, R., Weinberg, R.A., 2009. The basics of epithelial-mesenchymal transition. J. Clin. Invest. 119, 1420–1428. doi:10.1172/JCI39104

Karmodiya, K., Krebs, A.R., Oulad-Abdelghani, M., Kimura, H., Tora, L., 2012. H3K9 and H3K14 acetylation co-occur at many gene regulatory elements, while H3K14ac marks a subset of inactive inducible promoters in mouse embryonic stem cells. BMC Genomics 13, 424. doi:10.1073/pnas.0905443106

Kim, J.-W., Tchernyshyov, I., Semenza, G.L., Dang, C.V., 2006. HIF-1-mediated expression of pyruvate dehydrogenase kinase: a metabolic switch required for cellular adaptation to hypoxia. Cell Metabolism 3, 177–185. doi:10.1016/j.cmet.2006.02.002

Lee, H.-S., Park, J.-H., Kim, S.-J., Kwon, S.-J., Kwon, J., 2010. A cooperative activation loop among SWI/SNF, γ-H2AX and H3 acetylation for DNA double-strand break repair. EMBO J 1–12. doi:10.1038/emboj.2010.27

Love, M.I., Huber, W., Anders, S., 2014. Moderated estimation of fold change and dispersion for RNA-seq data with DESeq2. Genome Biol 15, 550. doi:10.1186/s13059-014-0550-8

Mahalingaiah, P.K.S., Ponnusamy, L., Singh, K.P., 2016. Oxidative stress-induced epigenetic changes associated with malignant transformation of human kidney epithelial cells. Oncotarget 5. doi:10.18632/oncotarget.12091

Mahalingaiah, P.K.S., Ponnusamy, L., Singh, K.P., 2015. Chronic oxidative stress leads to malignant transformation along with acquisition of stem cell characteristics, and epithelial to mesenchymal transition in human renal epithelial cells. J. Cell. Physiol. 230, 1916–1928. doi:10.1002/jcp.24922

Miao, D., Margolis, C.A., Gao, W., Voss, M.H., Li, W., Martini, D.J., Norton, C., Bossé, D., Wankowicz, S.M., Cullen, D., Horak, C., Wind-Rotolo, M., Tracy, A., Giannakis, M., Hodi, F.S., Drake, C.G., Ball, M.W., Allaf, M.E., Snyder, A., Hellmann, M.D., Ho, T., Motzer, R.J., Signoretti, S., Kaelin, W.G., Choueiri, T.K., Van Allen, E.M., 2018. Genomic correlates of response to immune checkpoint therapies in clear cell renal cell carcinoma. Science 373, eaan5951–1813. doi:10.1126/science.aan5951

Mo, D., Li, C., Liang, J., Shi, Q., Su, N., Luo, S., Zeng, T., Li, X., 2015. Low PBRM1 identifies tumor progression and poor prognosis in breast cancer. Int J Clin Exp Pathol 8, 9307–9313.

Murakami, A., Wang, L., Kalhorn, S., Schraml, P., Rathmell, W.K., Tan, A.C., Nemenoff, R., Stenmark, K., Jiang, B.-H., Reyland, M.E., Heasley, L., Hu, C.J., 2017. Context-dependent role for chromatin remodeling component PBRM1/BAF180 in clear cell renal cell carcinoma. Oncogenesis 2017 6:1 6, e287–e287. doi:10.1038/oncsis.2016.89

Nargund, A.M., Pham, C.G., Dong, Y., Wang, P.I., Osmangeyoglu, H.U., Xie, Y., Aras, O., Han, S., Oyama, T., Takeda, S., Ray, C.E., Dong, Z., Berge, M., Hakimi, A.A., Monette, S., Lekaye, C.L., Koutcher, J.A., Leslie, C.S., Creighton, C.J., Weinhold, N., Lee, W., Tickoo, S.K., Wang, Z., Cheng, E.H., Hsieh, J.J., 2017. The SWI/SNF Protein PBRM1 Restrains VHL-Loss-Driven Clear Cell Renal Cell Carcinoma. CellReports 18, 2893–2906. doi:10.1016/j.celrep.2017.02.074

Nogales-Cadenas, R., Carmona-Saez, P., Vazquez, M., Vicente, C., Yang, X., Tirado, F., Carazo, J.M., Pascual-Montano, A., 2009. GeneCodis: interpreting gene lists through enrichment analysis and integration of diverse biological information. Nucleic Acids Research 37, W317–W322. doi:10.1093/nar/gkp416

Pan, D., Kobayashi, A., Jiang, P., de Andrade, L.F., Tay, R.E., Luoma, A., Tsoucas, D., Qiu, X., Lim, K., Rao, P., Long, H.W., Yuan, G.-C., Doench, J., Brown, M., Liu, S., Wucherpfennig, K.W., 2018. A major chromatin regulator determines resistance of tumor cells to T cell–mediated killing. Science eaao1710. doi:10.1126/science.aao1710

Peña-Llopis, S., Vega-Rubín-de-Celis, S., Liao, A., Leng, N., Pavía-Jiménez, A., Wang, S., Yamasaki, T., Zhrebker, L., Sivanand, S., Spence, P., Kinch, L., Hambuch, T., Jain, S., Lotan, Y., Margulis, V., Sagalowsky, A.I., Summerour, P.B., Kabbani, W., Wong, S.W.W., Grishin, N., Laurent, M., Xie, X.-J., Haudenschild, C.D., Ross, M.T., Bentley, D.R., Kapur, P., Brugarolas, J., 2012. BAP1 loss defines a new class of renal cell carcinoma. Nat Genet 44, 751–759. doi:10.1038/ng.2323

Pfaffl, M.W., 2001. A new mathematical model for relative quantification in real-time RT-PCR. Nucleic Acids Research 29, e45.

Piva, F., Santoni, M., Matrana, M.R., Satti, S., Giulietti, M., Occhipinti, G., Massari, F., Cheng, L., Lopez-Beltran, A., Scarpelli, M., Principato, G., Cascinu, S., Montironi, R., 2015. BAP1, PBRM1 and SETD2 in clear-cell renal cell carcinoma: molecular diagnostics and possible targets for personalized therapies. Expert Review of Molecular Diagnostics 15, 1201–1210. doi:10.1586/14737159.2015.1068122

Porter, E.G., Dykhuizen, E.C., 2017. Individual Bromodomains of Polybromo-1 Contribute to Chromatin Association and Tumor Suppression in Clear Cell Renal Carcinoma. J Biol Chem 292, 2601–2610. doi:10.1074/jbc.M116.746875

RaMírez, F., Ryan, D.P., Grüning, B., Bhardwaj, V., Kilpert, F., Richter, A.S., Heyne, S., Dündar, F., Manke, T., 2016. deepTools2: a next generation web server for deep-sequencing data analysis. Nucleic Acids Research 44, W160–W165. doi:10.1093/nar/gkw257

Rawson, R.B., 2013. The site-2 protease. Biochimica et Biophysica Acta (BBA) - Biomembranes 1828, 2801–2807. doi:10.1016/j.bbamem.2013.03.031

Roupé, K.M., Veerla, S., Olson, J., Stone, E.L., Sørensen, O.E., Hedrick, S.M., Nizet, V., 2014. Transcription Factor Binding Site Analysis Identifies FOXO Transcription Factors as Regulators of the Cutaneous Wound Healing Process. PLoS ONE 9, e89274. doi:10.1371/journal.pone.0089274

Roy, N., Malik, S., Villanueva, K.E., Urano, A., Lu, X., Figura, von G., Seeley, E.S., Dawson, D.W., Collisson, E.A., Hebrok, M., 2015. Brg1 promotes both tumor-suppressive and oncogenic activities at distinct stages of pancreatic cancer formation. Gene Dev 29, 658–671. doi:10.1101/gad.256628.114

Rössler, O.G., Thiel, G., 2017. Specificity of Stress-Responsive Transcription Factors Nrf2, ATF4, and AP-1. J. Cell. Biochem. 118, 127–140. doi:10.1002/jcb.25619

Santner, S.J., Dawson, P.J., Tait, L., Soule, H.D., Eliason, J., Mohamed, A.N., Wolman, S.R., Heppner, G.H., Miller, F.R., 2001. Malignant MCF10CA1 cell lines derived from premalignant human breast epithelial MCF10AT cells. Breast Cancer Res Treat 65, 101–110.

Sato, Y., Yoshizato, T., Shiraishi, Y., Maekawa, S., Okuno, Y., Kamura, T., Shimamura, T., Sato-Otsubo, A., Nagae, G., Suzuki, H., Nagata, Y., Yoshida, K., Kon, A., Suzuki, Y., Chiba, K., Tanaka, H., Niida, A., Fujimoto, A., Tsunoda, T., Morikawa, T., Maeda, D., Kume, H., Sugano, S., Fukayama, M., Aburatani, H., Sanada, M., Miyano, S., Homma, Y., Ogawa, S., 2013. Integrated molecular analysis of clear-cell renal cell carcinoma. Nat Genet 45, 860–867. doi:10.1038/ng.2699

Schram, A.W., Baas, R., Jansen, P.W.T.C., Riss, A., Tora, L., Vermeulen, M., Timmers, H.T.M., 2013. A Dual Role for SAGA-Associated Factor 29 (SGF29) in ER Stress Survival by Coordination of Both Histone H3 Acetylation and Histone H3 Lysine-4 Trimethylation. PLoS ONE 8, e70035. doi:10.1371/journal.pone.0070035

Shain, A.H., Pollack, J.R., 2013. The spectrum of SWI/SNF mutations, ubiquitous in human cancers. PLoS ONE 8, e55119. doi:10.1371/journal.pone.0055119

Shen, C., Beroukhim, R., Schumacher, S.E., Zhou, J., Chang, M., Signoretti, S., Kaelin, W.G., 2011. Genetic and functional studies implicate HIF1α as a 14q kidney cancer suppressor gene. Cancer Discovery 1, 222–235. doi:10.1158/2159-8290.CD-11-0098

Shu, X.S., Zhao, Y., Sun, Y., Zhong, L., Cheng, Y., Zhang, Y., Ning, K., Tao, Q., Wang, Y., Ying, Y., 2017. PBRM1 restricts the basal activity of innate immune system through repressing RIG-I-like receptor signaling and is a potential prognostic biomarker for colon cancer. J. Pathol. doi:10.1002/path.4986

Soule, H.D., Maloney, T.M., Wolman, S.R., Peterson, W.D., Brenz, R., McGrath, C.M., Russo, J., Pauley, R.J., Jones, R.F., Brooks, S.C., 1990. Isolation and characterization of a spontaneously immortalized human breast epithelial cell line, MCF-10. Cancer research 50, 6075–6086.

Sun, X., Wang, S.C., Wei, Y., Luo, X., Jia, Y., Li, L., Gopal, P., Zhu, M., Nassour, I., Chuang, J.-C., Maples, T., Celen, C., Nguyen, L.H., Wu, L., Fu, S., Li, W., Hui, L., Tian, F., Ji, Y., Zhang, S., Sorouri, M., Hwang, T.H., Letzig, L., James, L., Wang, Z., Yopp, A.C., Singal, A.G., Zhu, H., 2017. Arid1a Has Context-Dependent Oncogenic and Tumor Suppressor Functions in Liver Cancer. Cancer Cell 32, 574–589.e6. doi:10.1016/j.ccell.2017.10.007

Suter, M.A., Chen, A., Burdine, M.S., Choudhury, M., Harris, R.A., Lane, R.H., Friedman, J.E., Grove, K.L., Tackett, A.J., Aagaard, K.M., 2012. A maternal high-fat diet modulates fetal SIRT1 histone and protein deacetylase activity in nonhuman primates. Faseb J 26, 5106–5114. doi:10.1096/fj.12-212878

Tatarskiy, V.V., Simonov, Y.P., Shcherbinin, D.S., Brechalov, A.V., Georgieva, S.G., Soshnikova, N.V., 2017. Stability of the PHF10 subunit of PBAF signature module is regulated by phosphorylation: role of β-TrCP. Sci. Rep. 7, 5645. doi:10.1038/s41598-017-05944-3

Tiwari, N., Meyer-Schaller, N., Arnold, P., Antoniadis, H., Pachkov, M., van Nimwegen, E., Christofori, G., 2013. Klf4 Is a Transcriptional Regulator of Genes Critical for EMT, Including Jnk1 (Mapk8). PLoS ONE 8, e57329.

Varela, I., Tarpey, P., Raine, K., Huang, D., Ong, C.K., Stephens, P., Davies, H., Jones, D., Lin, M.-L., Teague, J., Bignell, G., Butler, A., Cho, J., Dalgliesh, G.L., Galappaththige, D., Greenman, C., Hardy, C., Jia, M., Latimer, C., Lau, K.W., Marshall, J., Mclaren, S., Menzies, A., Mudie, L., Stebbings, L., Largaespada, D.A., Wessels, L.F.A., Richard, S., Kahnoski, R.J., Anema, J., Tuveson, D.A., Perez-Mancera, P.A., Mustonen, V., Fischer, A., Adams, D.J., Rust, A., Chan-On, W., Subimerb, C., Dykema, K., Furge, K., Campbell, P.J., Teh, B.T., Stratton, M.R., Futreal, P.A., 2011. Exome sequencing identifies frequent mutation of the SWI/SNF complex gene PBRM1 in renal carcinoma. Nature 469, 539–542. doi:10.1038/nature09639

Wang, Y., Kallgren, S.P., Reddy, B.D., Kuntz, K., López-Maury, L., Thompson, J., Watt, S., Ma, C., Hou, H., Shi, Y., Yates, J.R., III, Bähler, J., O’Connell, M.J., Jia, S., 2012. Histone H3 Lysine 14 Acetylation Is Required for Activation of a DNA Damage Checkpoint in Fission Yeast. The Journal of Biological Chemistry 287, 4386–4393. doi:10.1074/jbc.M111.329417

Wang, Y., Zhou, Y., Graves, D.T., 2014. FOXO Transcription Factors: Their Clinical Significance and Regulation. BioMed Research International 2014, 1–13. doi:10.1155/2014/925350

Wang, Z., Zhai, W., Richardson, J.A., Olson, E.N., Meneses, J.J., Firpo, M.T., Kang, C., Skarnes, W.C., Tjian, R., 2004. Polybromo protein BAF180 functions in mammalian cardiac chamber maturation. Gene Dev 18, 3106–3116. doi:10.1101/gad.1238104

Xia, W., Nagase, S., Montia, A.G., Kalachikov, S.M., Keniry, M., Su, T., Memeo, L., Hibshoosh, H., Parsons, R., 2008. BAF180 Is a Critical Regulator of p21 Induction and a Tumor Suppressor Mutated in Breast Cancer 68, 1667–1674. doi:10.1158/0008-5472.CAN-07-5276

Xu, S., Grullon, S., Ge, K., Peng, W., 2014. Spatial clustering for identification of ChIP-enriched regions (SICER) to map regions of histone methylation patterns in embryonic stem cells. Methods Mol. Biol. 1150, 97–111. doi:10.1007/978-1-4939-0512-6_5

Xue, Y., Canman, J.C., Lee, C.S., Nie, Z., Yang, D., Moreno, G.T., Young, M.K., Salmon, E.D., Wang, W., 2000. The human SWI/SNF-B chromatin-remodeling complex is related to yeast rsc and localizes at kinetochores of mitotic chromosomes. P Natl Acad Sci Usa 97, 13015–13020. doi:10.1073/pnas.240208597

Yu, T., Chen, X., Zhang, W., Li, J., Xu, R., Wang, T.C., Ai, W., Liu, C., 2012. Krüppel-like Factor 4 Regulates Intestinal Epithelial Cell Morphology and Polarity. PLoS ONE 7, e32492. doi:10.1371/journal.pone.0032492

